# Outpacing *E. coli*: Development of *Vibrio natriegens* as a Next-Generation Cloning Host

**DOI:** 10.64898/2026.02.11.705132

**Authors:** Eric Wei, Michael Louie, Emmett Dessimoz, Curtis Orona, Nicholas Smith, Natalie Holste, Michelle Slind, Han Nguyen, Santhosh Anandhan, Santhosh Kallivalappil, Matthew Weinstock

## Abstract

Despite transformative advances in DNA synthesis, sequencing, and automation that have accelerated recombinant DNA workflows, molecular cloning hosts have scarcely evolved past the *Escherichia coli* strains adopted out of convenience in the 1970s. We present NBx CyClone™ – an engineered strain of *Vibrio natriegens* – as a next-generation host for molecular cloning. This non-pathogenic marine bacterium combines broad plasmid and genetic tool compatibility, a versatile metabolism, and the fastest known doubling time of any free-living organism. By shortening growth-dependent steps, this host offers a practical route to faster, more efficient recombinant DNA workflows across research and industry.

## INTRODUCTION

Recombinant DNA technology is the foundation of modern biotechnology and is the essential starting point for any application in which biology is reprogrammed to serve human needs. From basic academic research to industrial bioproduction, our ability to manipulate DNA underpins a wide range of applications in the agricultural, consumer goods, therapeutic, environmental, materials, and energy sectors.

The foundational breakthroughs that made this possible occurred in the United States in the early 1970s. In 1972, Paul Berg created the first recombinant DNA molecule by fusing DNA from a bacteriophage with that of the SV40 virus^1^. Building on this, Herbert Boyer and Stanley Cohen developed the first practical method for generating and cloning recombinant DNA in 1973^2^. They used restriction enzymes to cut both a plasmid vector and a DNA fragment of interest, enzymatically joined them together with a ligase, and introduced the recombinant plasmid into *Escherichia coli*, where it was propagated. This achievement marked the birth of modern molecular cloning and genetic engineering. Shortly thereafter, Boyer co-founded Genentech, which achieved the first major commercial success with the technology via the production of recombinant human insulin, launching the biotech industry as we know it today^3^.

A central operation in molecular cloning workflows is the introduction of plasmid DNA into a bacterial host, a step which serves two main purposes. First, it provides a means to isolate a correct clone (i.e., DNA of the desired sequence). Methods for generating DNA fragments (either by PCR, chemical synthesis, or enzymatic synthesis) introduce errors into DNA at varying frequencies^4^. As most bacteria take up a single plasmid molecule and then faithfully replicate the DNA, clonal bacterial colonies isolated on agar plates can be screened to identify a colony containing the correct DNA sequence. Second, once a correct clone has been isolated, the bacterial host can be leveraged to generate a desired mass of plasmid DNA. As the bacteria replicates, each daughter cell will also inherit copies of the plasmid, allowing for the amplification of large DNA quantities. This can be scaled from small culture tubes in the laboratory (to generate microgram quantities of DNA), to large-scale industrial bioreactors (to generate kilograms of DNA for vaccine production).

While remarkable advances have been made since the time of Boyer and Cohen in terms of gene synthesis, DNA assembly, DNA sequencing, and automation (enabling high-throughput and large-scale manipulation of DNA) the microbial host used for cloning has remained largely unchanged. Pioneers of recombinant DNA technology chose *E. coli* primarily due to their familiarity with the organism, having been established as a model organism in bacteriology, genetics, and molecular biology for decades due to its ease of growth in the laboratory, well-characterized physiology, and ability to grow on inexpensive media. Today, most commercial molecular cloning workflows still rely on derivatives of the same *E. coli* strains developed decades ago with little attention to optimizing the host at the center of this process.

Considering advances in synthetic biology and in our understanding of microbial biodiversity, we contend that it is time to revisit the choice of cloning host. One particularly promising candidate for a next-generation molecular cloning host is *Vibrio natriegens*, a fast-growing, non-pathogenic, marine bacterium with several compelling features, including: compatibility with common plasmid systems used for research and industrial applications^5,6^, a rapidly expanding genetic toolkit^5,7–12^, a versatile metabolism^13–15^, and the fastest known doubling time of any free-living organism^16^.

This last feature is especially important. Time spent waiting for cloning hosts to grow (both as colonies on plates, and in liquid culture) constitutes a significant portion of the overall molecular cloning timeline. Accelerating these steps could yield meaningful acceleration in the design–build–test cycle that underpins synthetic biology and modern biotechnology.

In this manuscript, we describe the development of *Vibrio natriegens* as a next-generation host for molecular cloning, culminating in the engineering of a commercial strain – NBx CyClone™ – designed to accelerate molecular cloning and plasmid production workflows by reducing the time required for host growth.

## RESULTS

### Strain development

Modern *Escherichia coli* strains used for molecular cloning have diverged substantially from the progenitor isolates obtained from human fecal samples in the late 19^th^ and early 20^th^ centuries. Many of the genetic modifications were deliberately introduced to enhance their performance as cloning hosts, such as the removal of restriction systems or recombination machinery^17^. Others arose unintentionally through *E. coli’s* long history of laboratory cultivation, during which strains were subjected to repeated passaging, mutagenesis, and phage-mediated gene transfer^18–20^. Over time, these processes produced the highly adapted laboratory strains in use today, which are now so specialized for research settings that they have lost the ability to colonize the human gastrointestinal tract from which they originated^21,22^.

In contrast, *Vibrio natriegens* was first isolated in 1958 from salt marsh mud on Sapelo Island, Georgia, USA^16,23,24^ and subsequently preserved in various strain collections. Unlike *E. coli*, this organism has not been extensively passaged, remaining largely unchanged from its original environmental form and receiving little attention until recent years. In this study, we sought to introduce targeted genetic modifications into *V. natriegens* to improve its utility as a molecular cloning host.

#### Restriction systems

One of the most common genetic modifications in *E. coli* cloning strains is the removal of restriction-modification (R-M) systems. These systems act as bacterial defense mechanisms, protecting cells from bacteriophage infection by degrading foreign DNA; a process that can also interfere with the uptake and maintenance of recombinant plasmids. Therefore, for molecular cloning applications, R-M systems must be inactivated or eliminated. Bioinformatic analysis on the genome sequence of the type strain of *Vibrio natriegens* revealed that the strain lacks identifiable R-M systems, indicating that no such modifications are required to render it compatible with recombinant DNA.

#### Nucleases

Previous work^5^ demonstrated that deletion of the nuclease-encoding *dns* gene substantially improved the integrity of plasmid DNA isolated using commercial preparation kits by preventing DNA degradation by residual nucleases. In addition to enhancing plasmid quality, we found that the *dns* deletion also markedly increased transformation efficiency (Figure 1), which is a significant benefit for molecular cloning applications where transformation efficiency is a critical parameter.

**FIGURE 1:**
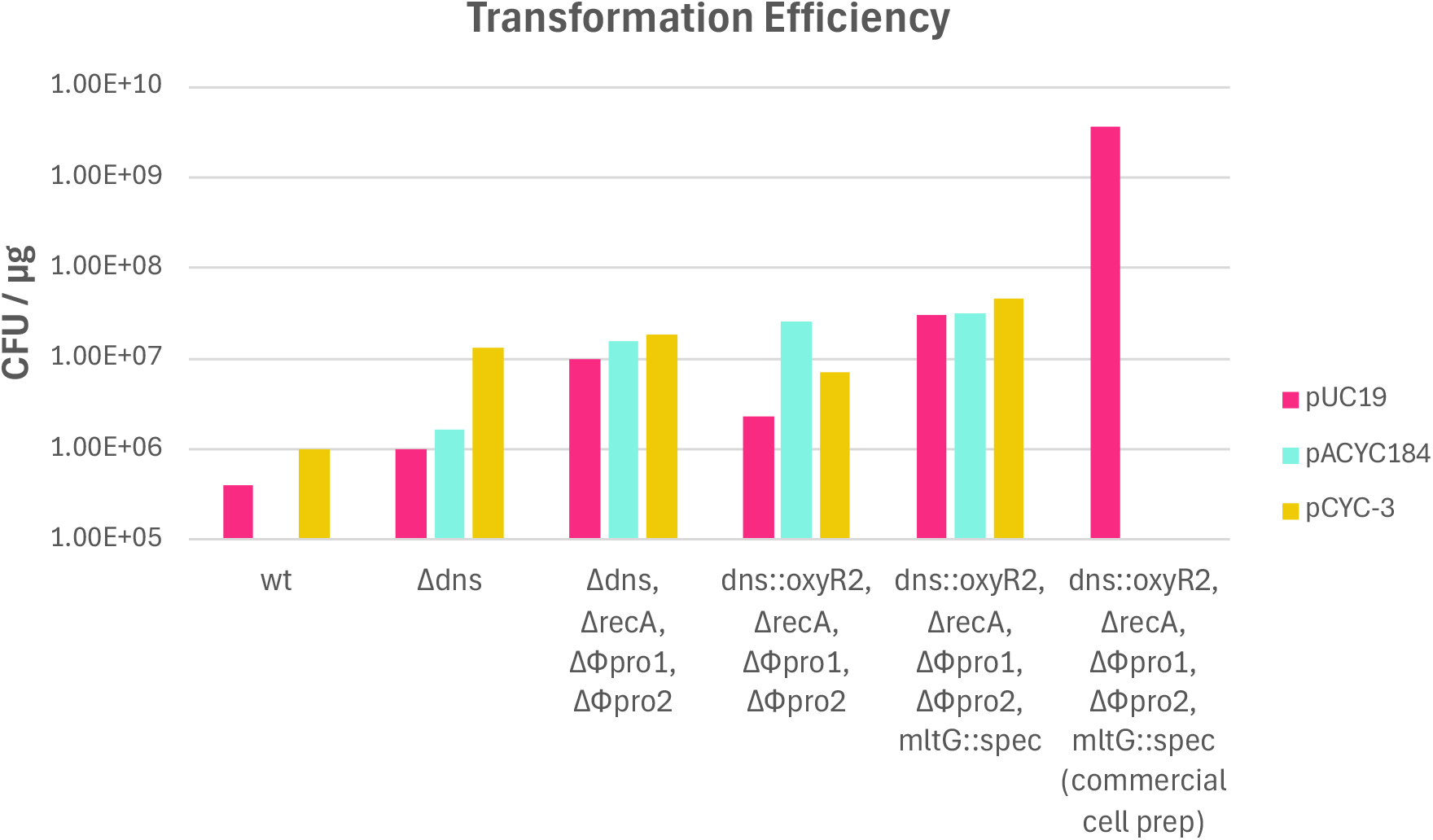
Transformation Efficiency of various engineered *V. natriegens* strains with select plasmids. The transformation efficiency of various engineered *V. natriegens* strains was assessed using plasmids pUC19 (agar with 50 mg/L carbenicillin), pACYC184 (agar with 5 mg/L chloramphenicol), and pCYC-3 (agar with 200 mg/L kanamycin). The x-axis shows which genetic modifications have been introduced into the strain with “wt” representing the type strain of the species. The right-most column shows the transformation efficiency of the final NBx CyClone™ strain using the commercial competent cell prep process (n=8) for pUC19 only. pCYC-3 contains a high-copy origin based on pUC19. The sequence of pCYC-3 is provided in the Supplementary Materials.

#### Recombination machinery

Most *E. coli* cloning strains are deficient for one or more recombination-related genes. In their native context, these genes are involved in the repair of damaged DNA; however, these functions can negatively impact the quality of plasmid DNA through undesirable recombination events leading to plasmid truncation, rearrangement, or concatenation. The most common mutation in *E. coli* cloning strains is the inactivation of the *recA* gene, which while making the host more susceptible to environmental stressors, significantly reduce plasmid recombination events^25,26^.

Upon observing concatenation in plasmids isolated from *V. natriegens*, we searched for homologous genes to *E. coli recA*, identifying a strong match on chromosome I (Supplementary Text 1). Removal of the *recA*-homolog markedly diminished concatenation events (Figure 2).

**FIGURE 2:**
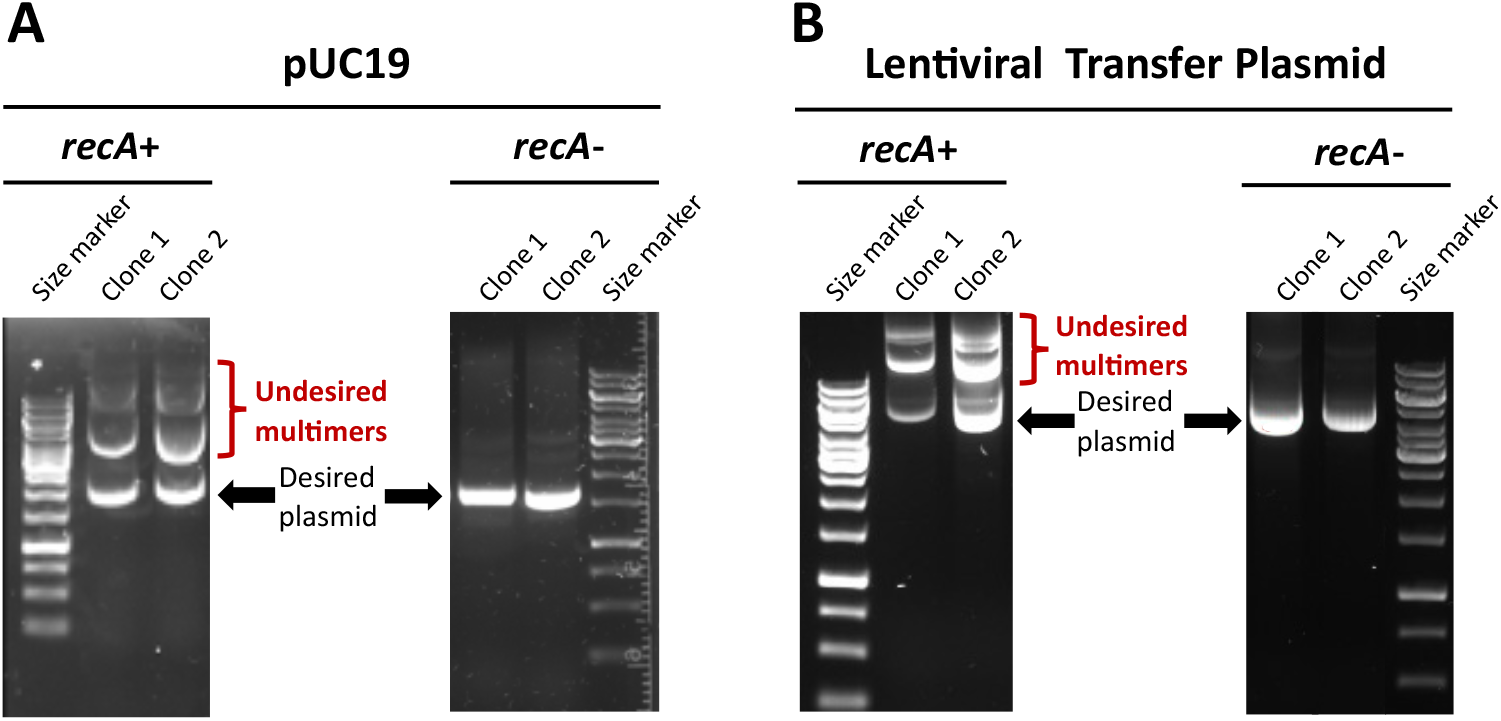
Topology of plasmid DNA isolated from *Vibrio natriegens* strains with and without *recA*. The commonly used pUC19 plasmid and a 7.4kb lentiviral transfer plasmid containing long terminal repeats (LTRs) were transformed into either a strain lacking only *dns* (***recA*+**), or a strain lacking both *dns* and *recA* (***recA*-**). Two independent colonies of each strain/plasmid combination were cultured in liquid medium followed by plasmid isolation via miniprep. Purified DNA samples were analyzed by agarose gel electrophoresis. The monomeric, supercoiled plasmid species is indicated as “Desired plasmid”, while higher-molecular-weight, concatenated species are indicated as “Undesired multimers”. In the case of both plasmids, the *recA* deletion leads to a marked reduction in plasmid concatenation.

#### Prophage

Bacterial genomes frequently harbor prophages – integrated viral genomes that remain dormant until triggered by environmental cues. Upon activation, these prophages can initiate phage production, which poses a significant risk in industrial and pharmaceutical manufacturing via catastrophic batch failures and costly facility contamination events^27^.

The *V. natriegens* type strain carries two prophage regions on chromosome I that have been shown to be inducible under standard laboratory conditions^28,29^. Notably, one report demonstrated that deletion of these two regions (referred to here as ϕpro1 and ϕpro2) conferred a strong fitness advantage, with the cured strain consistently outcompeting the wild-type strain under all tested osmotic conditions (hypo-osmotic, standard salt, and hyper-osmotic), and exhibited enhanced robustness against stressors such as osmotic imbalance and DNA damage^28^. To maximize strain stability and eliminate the risk of phage-related contamination, we removed both prophage regions from the genome, generating a Δϕpro1, Δϕpro2 strain.

#### Stress response genes

In contrast to laboratory strains of *E. coli* that benefit from over a century of adaptation, wild bacterial isolates are often sensitive to stressors unique to laboratory environments. One well-documented example is sensitivity to agar media. Many wild isolates exhibit impaired growth on agar plates due to low levels of peroxides in the agar generated during autoclaving^30,31^. This sensitivity has been observed in Vibrio species^32,33^, including *V. natriegens*^5^.

To address this limitation, researchers working with Vibrio species have supplemented agar with chemical compounds or enzymes that scavenge or degrade reactive oxygen species, or have engineered strains to overexpress heterologous catalase genes^5,32,34^.

Despite its peroxide sensitivity, *V. natriegens* encodes catalase homologs similar to those in *E. coli*, suggesting that the phenotype arises not from the absence of detoxification enzymes but from insufficient expression under laboratory conditions. Three reports specific to *V. natriegens* support this hypothesis. First, the organism shows weak *oxyR* (a conserved regulator of oxidative stress response genes) activation in the presence of peroxides, leading to low-level expression of catalase genes likely insufficient to provide a robust detoxification response^29^. Second, deletion of *oxyR* renders *V. natriegens* highly sensitive to oxidative stress^35^. Third, long-term cultivation under iron-replete conditions (which exacerbate oxidative stress), selects for mutations in *oxyR* (e.g., C208Y and C208G) that improve survival in oxidative environments and increase tolerance to peroxides^36^. Transcriptomic profiling of strains with these mutations revealed elevated constitutive expression of genes regulated by *oxyR*, including catalases, suggesting that increased expression underlies the protective effect.

Based on these observations, we engineered our strain to carry an additional copy of *oxyR* (bearing the C208G mutation) integrated at the *dns* locus (herein referred to as *oxyR2*). The *dns* locus was selected because it was previously shown to be a suitable locus for the integration of exogenous genes^5^. To assess whether this modification enhanced peroxide defense, we performed a catalase activity assay. As shown in Figure 3, the strain with only the wild-type copy of *oxyR* exhibited no visible bubbling upon exposure to hydrogen peroxide, whereas the strain harboring a copy of *oxyR* with the C208G allele displayed vigorous bubble formation, consistent with a stronger oxidative stress response.

**FIGURE 3:**
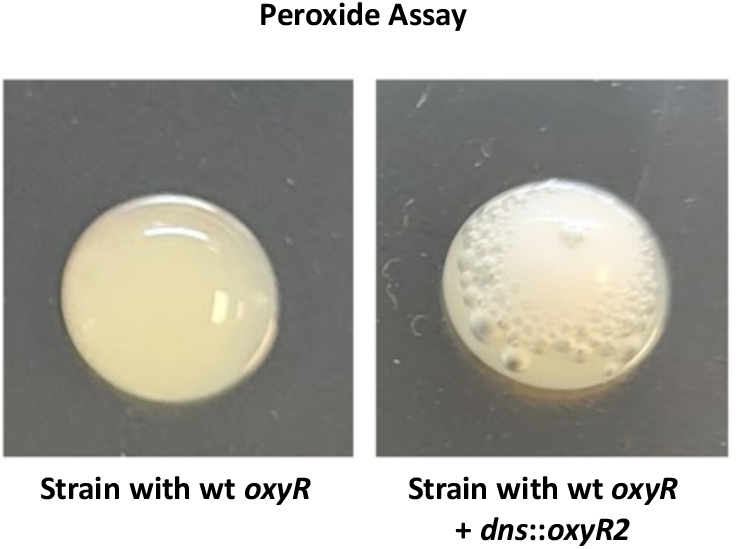
Peroxide assay performed on strains with and without *dns::oxyR2*. Liquid cultures of strains with and without the inclusion of *dns::oxyR2* (a second copy of *oxyR* carrying the C208G mutation inserted at the *dns* locus) were mixed with a hydrogen peroxide solution. The formation of bubbles is indicative of peroxide activity. As can be seen from the image, a strain carrying the wt copy of *oxyR* displays no evidence of peroxide activity (lack of bubbles), while the strain carrying an additional copy of *oxyR* with the C208G mutation shows a robust peroxide response.

We next tested the effect of this modification on plasmid DNA yields. As shown in Figure 4, the inclusion of *dns::oxyR2* increased plasmid yield by 60 – 80% after a six-hour cultivation. This effect was consistent across different media formulations, demonstrating that enhancing peroxide detoxification improves the robustness of *V. natriegens* for generalized laboratory applications including plasmid DNA production.

**FIGURE 4:**
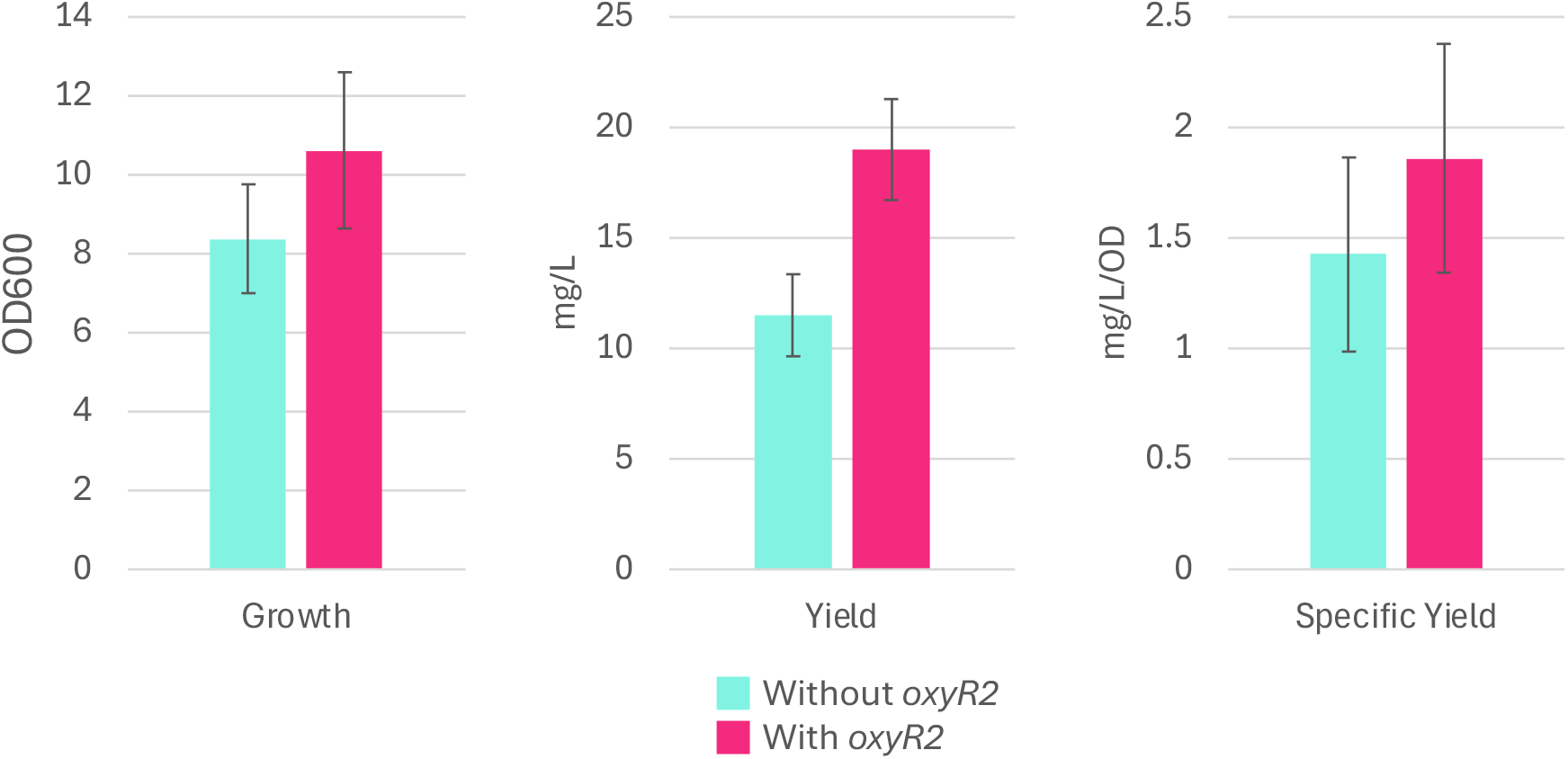
Plasmid yield and growth characteristics of strains with and without *dns::oxyR2*. Colonies of strains on agar plates with and without the *dns::oxyR2* modification carrying the plasmid pcDNA3.1 were inoculated into 2 mL aliquots of media in 24-well deep-well plates. Cultures were grown at 37°C with agitation at 300 RPM and humidity at 80% for 6 hours. Plasmid DNA was extracted from cell pellets from 100 μL of culture using Qiagen Miniprep columns, and plasmid DNA was quantified using a Qubit fluorometer. As seen from the figure, the presence of the *dns::oxyR2* genetic modification increased plasmid yield at the 6 hr timepoint.

#### Antibiotic resistance

*V. natriegens* is compatible with the standard suite of antibiotics used for molecular cloning applications, but its minimum inhibitory concentration (MIC) for kanamycin is substantially higher than that of *E. coli* (Supplementary Text 2). This natural tolerance necessitates the use of elevated kanamycin concentrations and increases the risk of background colony growth. We pursued multiple genetic strategies to reduce this natural resistance.

One well-described resistance mechanism against aminoglycosides (such as kanamycin) is active efflux, mediated by genetically encoded multidrug efflux pumps^37^. To test whether this contributed to resistance in *V. natriegens*, we bioinformatically identified a comprehensive list of predicted efflux pump genes and generated individual knockout strains (Supplementary Text 3). None of these mutants exhibited significantly increased kanamycin sensitivity (Supplementary Text 3). Although functional redundancy between pumps could obscure effects of single deletions, we did not pursue higher-order knockouts.

We also analyzed a published RNA-seq dataset from *V. natriegens* cultivated in M9 medium in the presence or absence of subinhibitory concentrations of kanamycin^38^, reasoning that transcriptional profiling might reveal genes upregulated in response to kanamycin that contribute to resistance. Differential expression analysis highlighted several candidates (Supplementary Text 4), but singlet deletion of each of the selected genes failed to significantly impact sensitivity (Supplementary Text 4).

As a final approach, we investigated whether alterations to the cell envelope might impact kanamycin susceptibility. Vibrio species, including *V. natreigens*^8,39^, are known to produce exopolysaccharides involved in biofilm formation, which have been reported to increase antibiotic tolerance in bacteria^40^. In a related manner, several reports indicate that mutations affecting peptidoglycan homeostasis influence antibiotic susceptibility. In particular, deletion of *mltG* (a lytic transglycosylase involved in glycan strand termination) has been shown to alter cell wall structure and alter antibiotic sensitivity in diverse Gram-negative species, including *Neisseria gonorrhoeae*^41^ and *Pseudomonas aeruginosa*^42,43^.

To test these possibilities, we constructed *V. natriegens* strains lacking capsular and exopolysaccharide genes/clusters, or the *mltG* homolog, and evaluated their antibiotic susceptibility. None of the knockouts displayed a significant reduction in kanamycin resistance (Supplementary Text 5). However, the *mltG* mutant reproducibly exhibited increased plasmid transformation efficiency (Figure 1), and this knockout was therefore retained in our final engineered strain, which we refer to as NBx CyClone™. NBx CyClone™ competent cells were able to achieve chemical transformation efficiencies exceeding 1.0E9 CFU/μg of plasmid DNA and maintain competence for at least 6 months (longest timepoint tested) when stored at -80°C (Supplementary Text 6).

### Media development

In parallel with development of the NBx CyClone™ strain, we sought to formulate growth media optimized for plasmid DNA production. Our media design criteria emphasized low cost, absence of animal-derived components, and the ability to support both rapid growth and high plasmid yield.

Previous studies showed that *V. natriegens* grows in standard broths used for *E. coli* (e.g., LB, BHI), with performance enhanced by elevated salt concentrations, leading to the widely used LBv2 recipe (based on LB Miller supplemented with MgCl_2_, KCl, and additional NaCl^5^).

To meet our design goals, we selected animal-free tryptone and yeast extract as nutrient sources, supplemented with MgSO_4_, NaCl, and K_2_HPO_4_to provide the most abundant marine ions (*V. natriegens* is a marine organism). Although the phosphate anion is not a dominant counterion in seawater, it was included for its buffering capacity and to ensure DNA synthesis was not phosphate limited. Early experiments confirmed that culture pH increased significantly during growth in rich media, likely due to ammonia release from amino acid catabolism. Increasing the phosphate concentration to add more buffering capacity resulted in issues with precipitation due to the interaction between magnesium and phosphate ions. We therefore evaluated the incorporation of additional buffering agents, finally settling on Tris (see Supplementary Text 7).

Having defined our components, we then applied a design-of-experiments (DOE) approach to optimize component concentrations, using plasmid yield after 4 hours of growth as the primary response variable (Supplementary Text 8). Solid agar media was developed using a similar approach (Supplementary Text 9), this time using transformation efficiency and colony growth as the response variables. These optimized medias were used for subsequent studies unless otherwise specified.

#### Acceleration of recombinant DNA workflows

With optimized strain and media in hand, we set out to evaluate if the rapid growth rate of *V. natriegens* could accelerate recombinant DNA workflows. We transformed NBx CyClone™, *E. coli* DH5α, and *E. coli* NEB® Turbo each with the high-copy mammalian expression plasmid pcDNA3.1. DH5α served as a benchmark strain due to its widespread use as for molecular cloning and plasmid production, whereas NEB® Turbo was included as a commercially available *E. coli* strain marketed for accelerating recombinant DNA workflows due to rapid growth rate. Transformed colonies were used to inoculate 2 mL cultures and cultivated for 6 hours before harvest. Growth kinetics, plasmid yield, and plasmid quality were then assessed for each strain (Figure 5).

**FIGURE 5:**
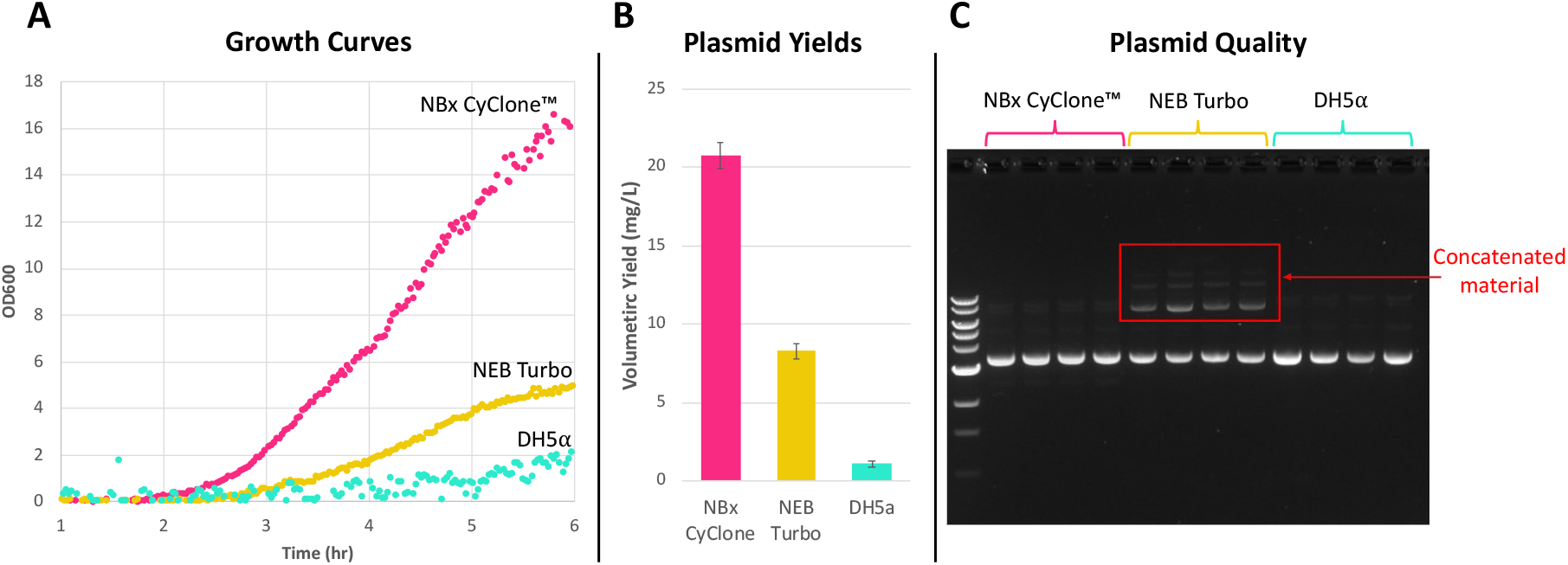
Comparison of NBx CyClone to DH5α and NEB® Turbo in accelerated plasmid production workflow. All strains were transformed with pcDNA3.1 via chemical transformation and plated on agar plates (NBx CyClone™ agar for NBx CyClone™, and LB for *E. coli*). Agar plates were incubated overnight, followed by colony picking into 2 mL aliquots of liquid media with carbenicillin in wells of a 24-well deep-well plate (In this preliminary experiment, DOE condition 14 (see Supplementary Text 8) media was used for NBx CyClone™, LB for *E. coli*). The plate was incubated at 37°C for 6 hours with agitation at 300 RPM in a shaker with a 25 mm orbit. **(A)** Growth curves measured directly in the 24-well plate via an ESSItech mtpOD online biomass monitor. **(B)** Average volumetric yields (in mg/L) of pcDNA3.1 after 6 hours of cultivation isolated by Qiagen miniprep columns and measured via fluorometry (Qubit). Averages were calculated from yields from four independent colonies (n=4) from each strain. **(C)** Plasmid quality assessed by a supercoiled agarose gel stained with SYBRsafe stain. The supercoiled gel allows visualization of other plasmid topologies such as undesired concatenated multimers.

As shown in Figure 5, panel A, the growth rate of *V. natriegens* substantially exceeded that of both *E. coli* strains, including the fast-growing NEB® Turbo. Volumetric plasmid yields (mg/L, Figure 5, panel B) mirrored the growth trends, with NBx CyClone™ producing the highest yield, NEB® Turbo showing an intermediate yield, and DH5α the lowest.

Plasmid topology was also assessed, as recombination events within the host can lead to undesirable truncations, rearrangements, or plasmid concatenation (multimer formation). Large-scale recombination events (such as large deletions or concatenation) are readily detected by supercoiled agarose gel electrophoresis, in which plasmid DNA is analyzed on an agarose gel without prior restriction digestion to preserve topological differences.

The supercoiled gel (Figure 5, panel C) revealed that plasmids isolated from NBx CyClone and *E. coli* DH5α were predominantly monomeric and supercoiled, whereas plasmid from NEB® Turbo contained an appreciable percentage of material in higher-order concatenated form. This finding is consistent with the *recA+* genotype of NEB® Turbo, which enhances growth by restoring DNA repair capacity (partially explaining the faster growth rate compared to other *E. coli* strains), but at the cost of increasing plasmid recombination events (which is why *recA* was removed from most modern cloning strains).

To further evaluate the performance of our new strain, 15 additional plasmids representing different antibiotic resistances, origins of replication, sizes, and downstream applications were transformed into NBx CyClone™ and DH5α (summarized in Supplementary Text 11). Transformation reactions were plated out on agar plates and incubated overnight. The following morning colonies were inoculated into 2 mL cultures and cultivated in a shaking incubator for 6 hours before harvest. At this point, all NBx CyClone™ cultures showed significant growth, while only a few *E. coli* DH5α cultures showed any measurable growth (Figure 6).

**FIGURE 6:**
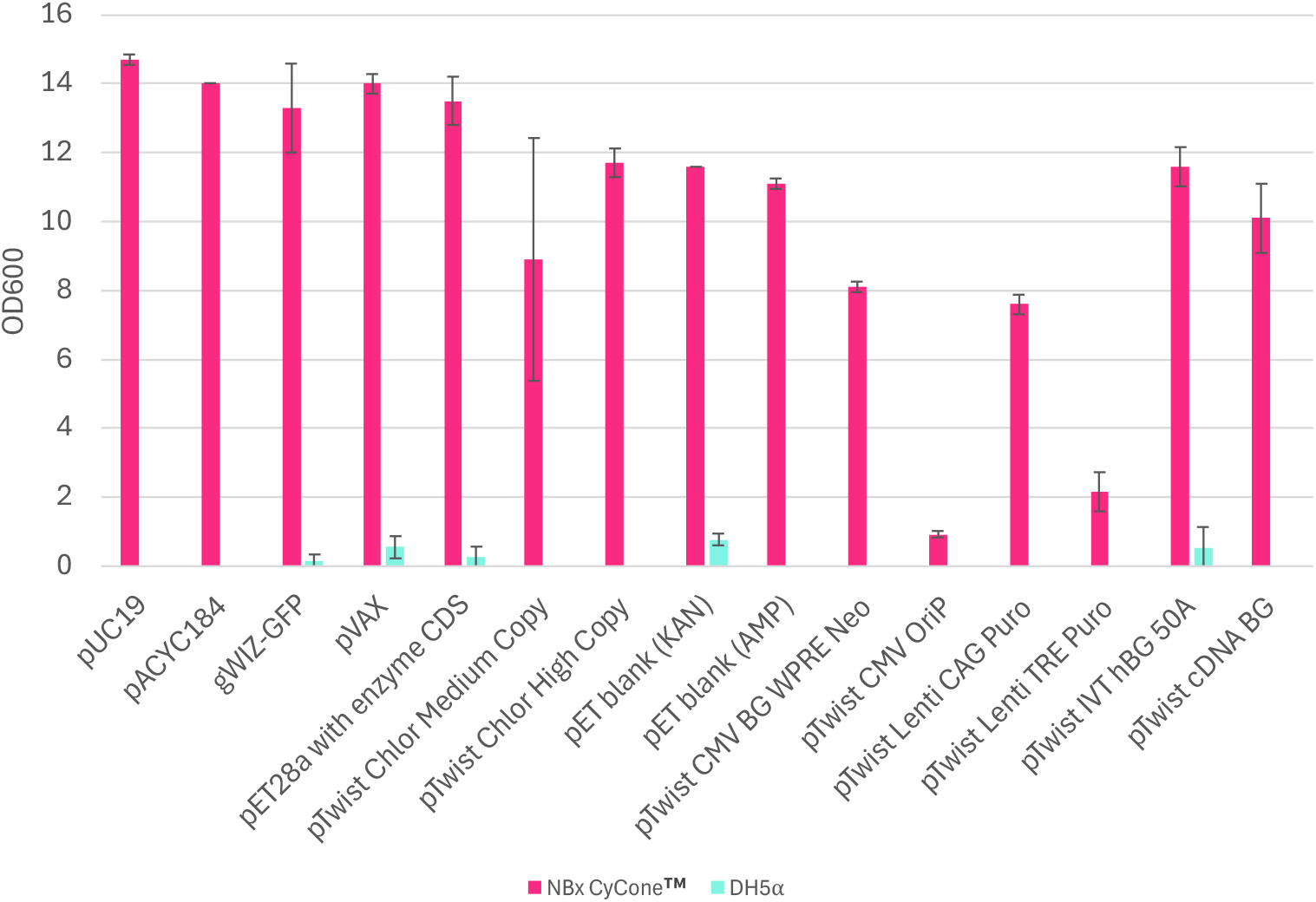
Growth of NBx CyClone and DH5α strains harboring various plasmids after 6 hours of growth. NBx CyClone™ or DH5α strains carrying a variety of common plasmids were streaked out on agar plates and incubated overnight at 37°C. After ∼16 hours of incubation, single colonies were inoculated into 2 mL of media (LB for *E. coli*, NBx CyClone™ growth media for NBx CyClone™) in 24-well deep-well plates and cultivated in a shaking incubator at 37°C for 6 hours. Biomass was measured spectroscopically by determining OD600. As can be observed in the graph, there was measurable growth for all 15 NBx CyClone™ cultures, but only modest growth in 5 *E. coli* cultures, with 10 showing no growth at the 6-hour timepoint.

Due to the lack of biomass from the *E. coli* cultivations, only the NBx CyClone™ samples were moved forward for plasmid extraction (along with additional plasmids that were not tested in *E. coli*). As can be seen from Figure 7, high yields can be generated in this shortened timeframe, saving considerable time in a typical design-build-test cycle.

**FIGURE 7:**
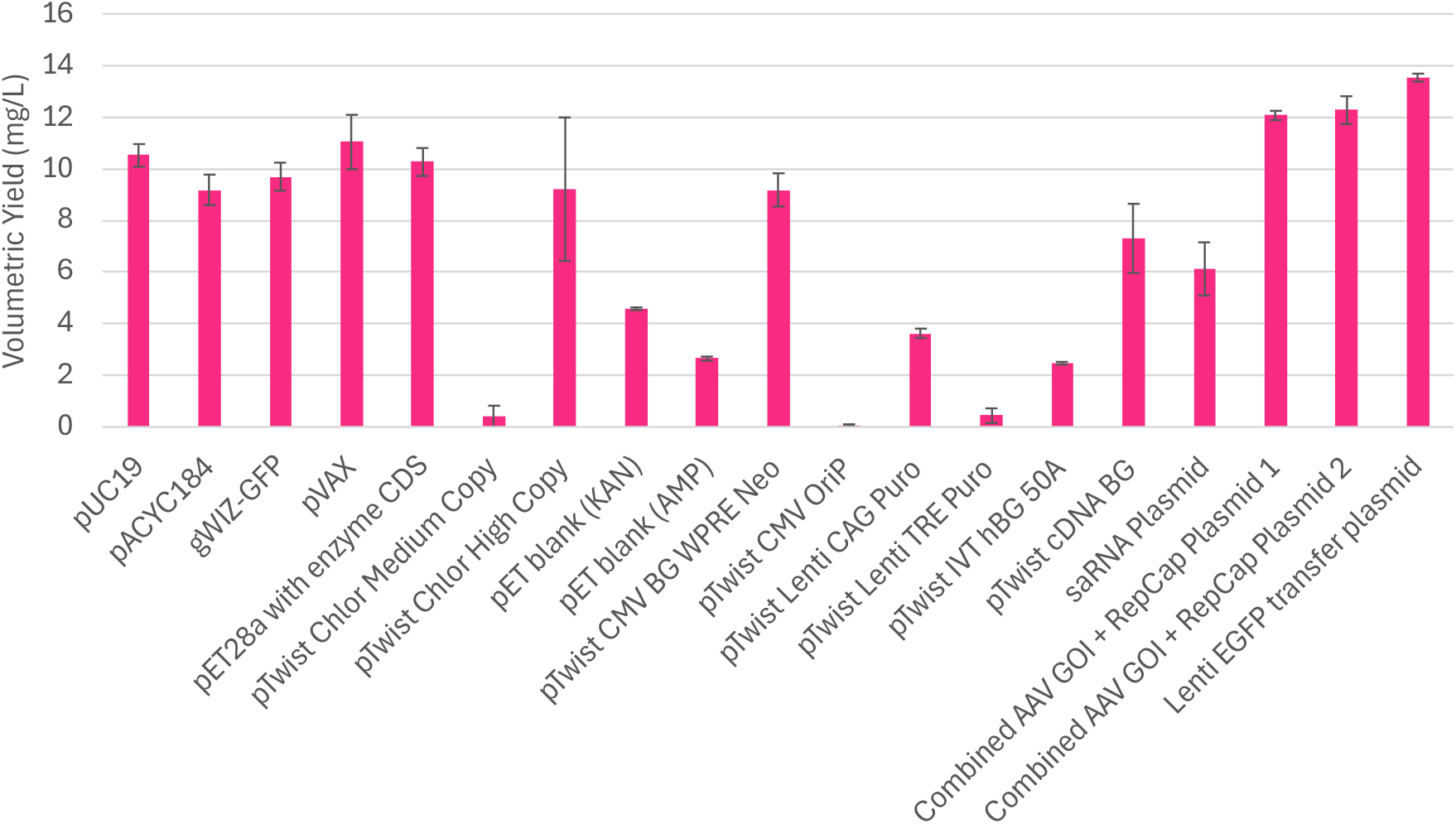
Plasmid yields from NBx CyClone™ strains after 6 hours of growth. Colonies of NBx CyClone™ strains harboring various plasmids were inoculated into 2 mL of media (NBx CyClone™ growth media) in 24-well deep-well plates and cultivated in a shaking incubator at 250 – 300 RPM at 37°C for 6 hours. Volumetric yields determined by harvesting 200 μL aliquots of each culture and processing through a Qiagen QIAprep® miniprep column. Plasmid concentrations were measured using a Qubit fluorometer.

To explore the flexibility of cultivation format, we also examined plasmid yields from cultures at 4, 6, and 8 hours in 96-well, 24-well, and 125 mL shake flask formats, each inoculated from a single NBx CyClone™ colony (Figure 8). Even the smallest culture volumes (0.5 – 4 mL) produced several micrograms of plasmid in 4 hours (sufficient for standard miniprep workflows), with yields increasing as culture time is extended. Larger-volume cultures (25 mL) achieved midiprep-scale yields after 6-8 hours, demonstrated that NBx CyClone™ supports rapid, scalable plasmid production across common laboratory formats.

**FIGURE 8:**
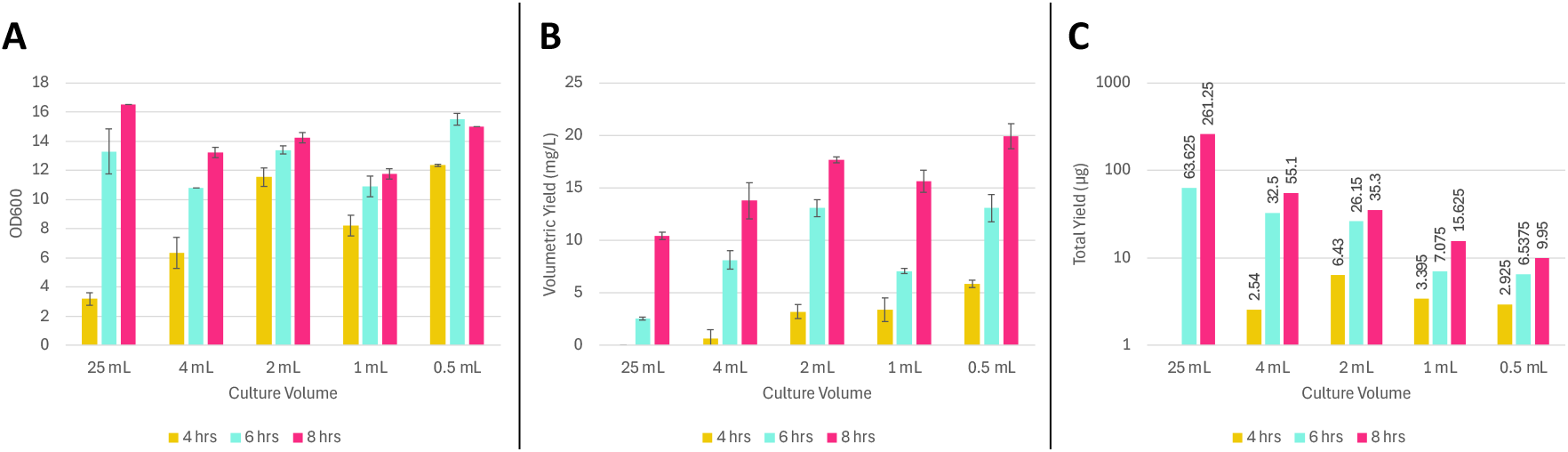
Growth and plasmid yield with NBx CyClone™ cultivated in different culture formats and incubation times. Individual colonies of NBx CyClone™ transformed with pUC19 were inoculated into different growth formats (25 mL culture volume in 125 mL baffled flask; 4 mL or 2 mL in a 24-well deep-well plate; and 1 mL or 0.5 mL in a 96-well deep-well plate). Each condition was tested in triplicate. All cultures were incubated at 37°C for 4, 6, or 8 hours. The flasks were agitated at 250 RPM with a 2.5 cm orbit, the 24-well deep-well plate at 300 RPM with a 2.5 cm orbit, and the 96-well deep-well plate at 1000 RPM with a 3 mm orbit. **(A)** Culture density (OD600) of each culture at different timepoints. **(B)** Volumetric yields (mg/L) determined by harvesting 200 μL aliquots of each culture and processing using a Qiagen QIAprep® miniprep kit. Plasmid concentrations were measured using a Qubit fluorometer. **(C)** Total plasmid (in µg) recovered from the respective culture volumes at 4, 6, and 8 hours.

#### Molecular Cloning

Having established that NBx CyClone™ can accelerate the growth-related steps in recombinant DNA workflows, we shifted our attention to evaluating the compatibility of NBx CyClone™ with common molecular cloning workflows. We performed plasmid assembly reactions using Gibson Assembly^44^, Golden Gate Assembly^45^, and Ligation-independent Cloning (LIC)^46^, and transformed the assemblies into NBx CyClone™ by chemical transformation. In all cases, transformations resulted in significant colonies (see Gibson Assembly data in Figure 9 and LIC data in Supplementary Text 10), suitable for routine cloning and library construction applications. Plasmid isolation and analysis confirmed that the recovered constructs matched the expected assemblies, demonstrating that NBx CyClone™ supports efficient and reliable cloning workflows.

**FIGURE 9:**
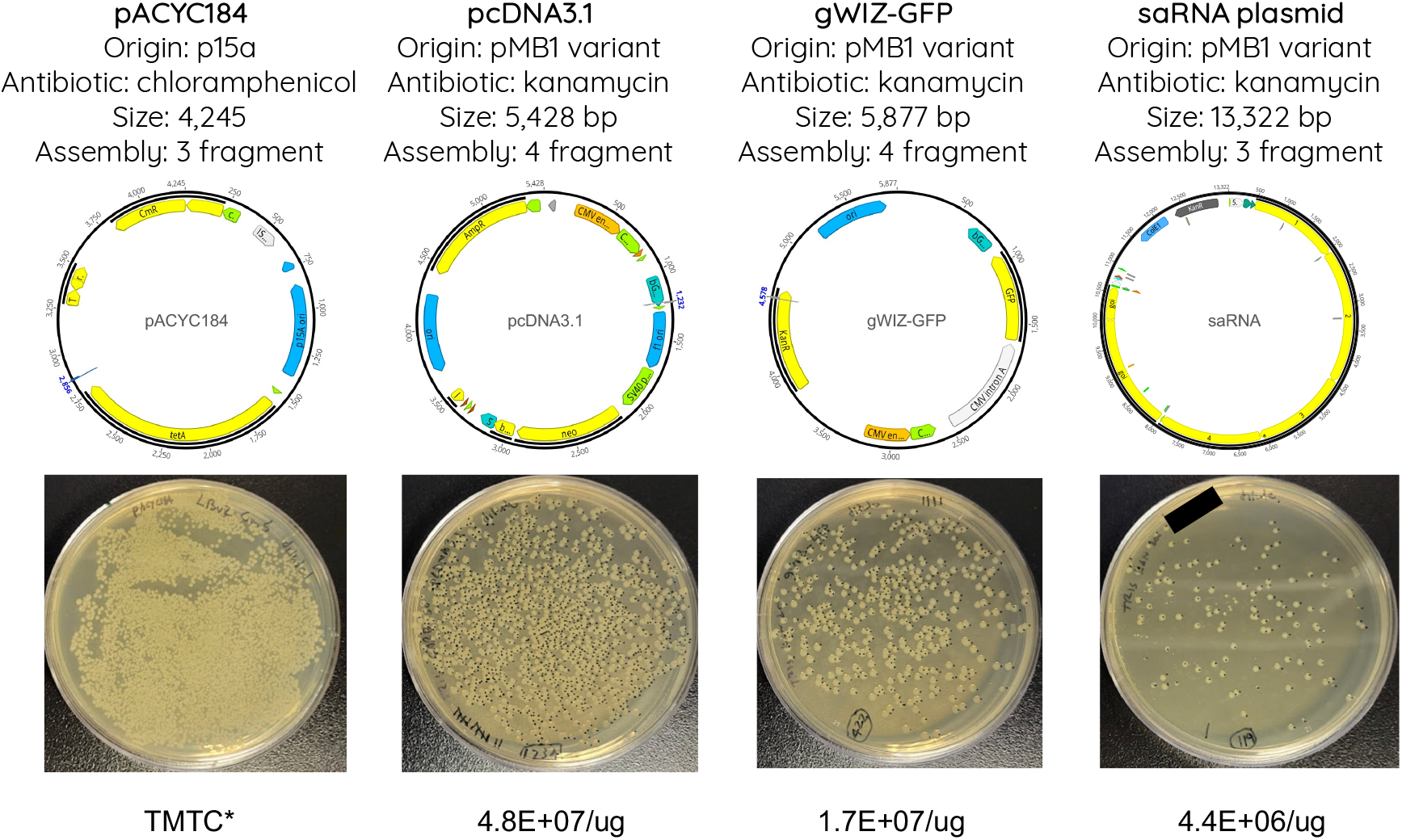
Representative data from transformations of NBx CyClone™ with Gibson Assembly reactions. Three common commercial plasmids (pACYC184, pcDNA3.1, gWIZ-GFP) and one internal self-amplifying RNA construct (saRNA) were broken into 3 – 4 fragments with overlapping homology by PCR. Gibson Assembly reactions were performed according to the manufacturer’s instructions (NEBuilder® HiFi DNA Assembly Master Mix) except that reaction volumes were reduced to 10 µL, and 0.025 pmol of each fragment were used in the assembly. 1 µL of each assembly reaction was used for transformation reactions. Post transformation recovery, 3 µL of recovered cells were diluted in 97 µL of fresh recovery media and the whole volume was plated out on agar plates. Plates were incubated overnight at 37°C. Colonies were enumerated and used to calculate transformation efficiencies. TMTC denotes “too many to count”.

#### Cloning plasmid backbones

To facilitate efficient routine cloning in NBx CyClone™, we designed a set of standardized high-copy cloning vectors collectively referred to as the pCYC series. These vectors were developed to provide flexibility in selection and assembly while maintaining compatibility with modern cloning workflows. Each vector incorporates a distinct antibiotic resistance marker to enable parallel cloning applications and features a high-copy pUC origin of replication to support robust plasmid yields. To ensure seamless Golden Gate compatibility, all internal BsaI recognition sites were eliminated, and a unique BsaI-based entry site was introduced to allow rapid and directional DNA assembly. The BsaI sites can also be used for rapid linearization of the backbone for use with Gibson Assembly workflows. The overall architecture of these plasmids is shown in Figure 10 and the sequence details are provided in Supplementary Text 12.

**FIGURE 10:**
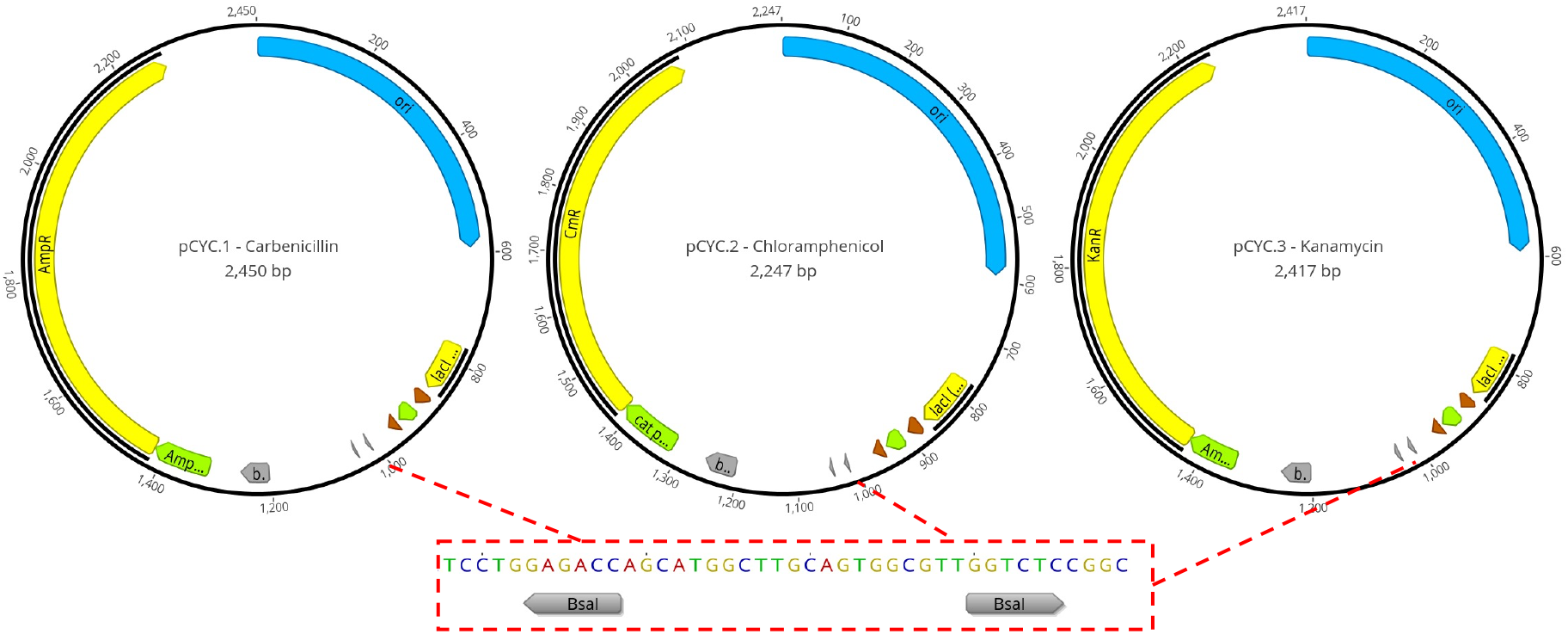
pCYC series cloning vectors. Three high-copy cloning vectors were designed for routine cloning applications in NBx CyClone™. Key features of each plasmid include a high-copy origin-of-replication, an antibiotic resistance marker (AmpR, CmR, and KanR varieties), removal of all instances of the BsaI restriction site from the backbone, and a unique BsaI entry site to facilitate rapid cloning by Golden Gate Assembly, or convenient plasmid linearization for Gibson Assembly or traditional restriction/ligation workflows.

## CONCLUSION

NBx CyClone™ represents a next-generation host for molecular cloning and plasmid DNA production. Through targeted genetic engineering we generated a strain that combines exceptional growth rate, genetic stability, and compatibility with standard molecular biology tools. When paired with optimized media formulations, NBx CyClone™ consistently outperformed benchmark *E. coli* strains, achieving higher plasmid yields, and markedly faster cultivation times.

By shortening the growth-dependent steps that dominate the molecular cloning workflow, NBx CyClone™ offers a practical and immediately deployable route to accelerate design–build–test cycles across basic research and industrial biotechnology, an outcome that has broad implications for both laboratory productivity and commercial DNA manufacturing.

More broadly, this work underscores the value of re-examining long-standing microbial tools through the lens of modern synthetic biology. Just as *E. coli* enabled the first era of recombinant DNA technology, *Vibrio natriegens*, with its exceptional growth rate and emerging genetic toolkit, is poised to power the next.

## MATERIALS AND METHODS

### Bacterial strains and cultivation

The type strain of *V. natriegens* was used as the starting material for this work (deposited in various strain collections as DSM 759, ATCC 14048, NCMB 857, NBRC 15636). The most common liquid medias for cultivation of the strain were LBv2^5^ (good for generic growth, but not plasmid production), and NBx CyClone™ Growth Media (optimized for rapid plasmid production), which is composed of 15 g/L yeast extract, 2.5 g/L vegetable tryptone, 4 g/L magnesium sulfate (anhydrous), 22.5 g/L sodium chloride, 3 g/L dipotassium phosphate, 150 mM Tris, pH 6.4. Media was typically sterilized via 0.2 micron filtration instead of autoclaving. Cultivations were performed in a variety of different formats: 0.5 – 1 mL cultures in 96-well deep-well plates incubated at 37°C with agitation at 900-1000 RPM with a 3 mm orbit and humidity at 80%; 2 – 4 mL cultures in 24-well deep-well plates incubated at 37°C with agitation at 300 RPM with a 2.5 cm orbit and humidity at 80%; 25 – 500 mL cultures in disposable baffled flasks (Thomson Ultra Yield®) sized 5x larger than the culture volume (e.g., a 25 mL culture is grown in a 125 mL flask) and incubated at 37°C with agitation at 250-300 RPM with a 2.5 cm orbit and humidity at 80%. The most common agar medias used for cultivation were LB Miller agar, LBv2 agar (LBv2 media with 1.5% agar), or NBx CyClone™ agar, which is composed of 15 g/L agar, 5 g/L yeast extract, 10 g/L vegetable tryptone, 19.54 g/L sodium chloride, 3.82 g/L magnesium sulfate (anhydrous), 117 mM Tris (from 1 M, pH 7 stock). Agar plates are made by combining the ingredients, mixing, autoclaving (121°C for 20 minutes on liquid cycle), cooling on stir plate until ∼55°C, adding antibiotic, pouring 20-25 mL plates (100 mm dishes) in a biosafety cabinet, allowing plates to solidify for ∼30 minutes, and then storing at 4°C. The following antibiotics were used in both liquid and agar media: Kanamycin (200 – 250 µg/mL), Carbenicillin (50 µg/mL), Chloramphenicol (5 µg/mL), Spectinomycin (200µg/mL), and Tetracycline (15 µg/mL). Agar plates were incubated between 30°C - 37°C, with 37°C being most typical.

### Plasmids

Test plasmids used in this work are summarized in Supplementary Text 11 and 12.

### Strain engineering

Gene knockouts were performed by replacing the target gene with a chloramphenicol or spectinomycin antibiotic resistance cassette using natural competence-mediated transformation as described in Des Soye et al.^47^. Successful knockouts were confirmed by colony PCR. In some cases, the antibiotic cassette was flanked on both sides by engineered loxP sites, allowing for the removal of the marker from the genome via Cre-lox recombination as described in Weinstock et al.^5^. Removal of the antibiotic marker was confirmed first by patch plating on agar plates with and without antibiotic, followed by colony PCR, and finally by extracting genomic DNA (Zymo Quick-DNA Fungal/Bacterial Miniprep Kit) followed by full genome sequencing (Plasmidsaurus).

### Comp cell prep & transformations

Competent cells were produced at Novel Bio and are available commercially as NBx Cyclone™. Transformations were performed as follows: one vial of NBx CyClone™ competent cells was thawed on ice for 30 minutes. The tube was gently flicked to mix contents and then 50 µL of cells was gently transferred to a pre-chilled 1.5 mL microcentrifuge tube. 1-2 µL of plasmid DNA was added to the cells and gently flicked to mix. Cells were incubated on ice for 20 minutes, heat shocked in a water bath at 42°C for 45 seconds, then returned to ice for 3 min. Cells were recovered by adding 1 mL of room temperature NBx CyClone™ Recovery Media and incubating at 37°C with agitation at 250-300 RPM for 1-2 hrs. Appropriate dilutions were made in fresh NBx CyClone™ Recovery Media, followed by plating 50 – 200 µL on pre-warmed agar plates. Plates were incubated between 6 – 16 hrs.

### Plasmid yield & quality

Plasmid yield and quality were assessed by extracting DNA via miniprep. Typically, cell pellets were harvested from 100 – 200 µL of culture by centrifugation at 5000 x g for 5 min, followed by removal of the supernatant. Cells were processed through a Qiagen QIAprep Spin Miniprep column following the manufacturer instructions with the following modifications: the optional PB wash step was used, and plasmid was eluted by 2 x 50 µL elutions with TE buffer to ensure complete recovery of plasmid from the column matrix. Plasmid volumetric yield was assessed by measuring the concentration of the recovered plasmid DNA on a Qubit™ fluorometer using the 1x dsDNA Broad Range Assay Kit, and then normalizing to the amount of culture processed through the column. Plasmids were sequenced by long-read NGS (ONT at Plasmidsaurus). Supercoiled gels were run by mixing purified plasmid DNA with 5x gel loading dye (NEB), loading on a hand-poured 1% agarose TAE gel containing SYBR Safe gel stain (Thermo Fisher), and running the gel at 120V for > 1 hr. Gels were visualized in a gel imager equipped with blue light illumination.

### Catalase assay

50 µL of a liquid culture of bacterial cells were pipetted onto a plastic surface. 10 μL of 3% Hydrogen peroxide was added to the sample and mixed by pipetting. After about 20 seconds, the samples were imaged. The presence of bubbles is indicative of catalase activity (in breaking down the hydrogen peroxide into water and gaseous oxygen).

### RNA seq analysis

The Geneious Prime software package was used to map RNA-seq reads to the *V. natriegens* chromosomes (accession numbers CP016345 and CP016346) and calculate expression levels. Expression levels between different cultivation conditions were compared within Geneious Prime using DESeq2. Data was visualized directly in Geneious Prime or exported to CSV for analysis. See Supplementary Text 4 for more details.

### DOE methodology

Details of the use of DOE methodology to aid in optimizing liquid and agar media formulations are included in the Supplementary Materials.

### Plasmid assembly

Gibson assemblies were performed in PCR tubes by mixing 0.025 pmol of each fragment in 5 µL total volume of molecular biology grade water and then adding 5 µL of 2x HiFi NEBuilder Master Mix (NEB). Reactions were incubated at 50°C in a thermocycler for 30 – 60 min. Golden Gate assemblies using BsaI sites were performed with the NEBridge® Golden Gate Assembly Kit BsaI-HF®v2 (NEB #E1601). Reactions (20 µL total) contained 1 µL backbone plasmid normalized to 10 ng/kbp, insert DNA (from plasmids or purified PCR amplicons) at 30 ng/kbp to achieve a 3:1 insert-to-backbone molar ratio of fragments, 2 µL T4 DNA Ligase Buffer, 2 µL Golden Gate Assembly Enzyme Mix, and nuclease-free water. Assemblies were cycled 30 times between 37 °C for 1 min and 16 °C for 1 min, followed by 60 °C for 5 min and a 4 °C hold. 3 µL of completed reactions were used in transformations. LIC reactions were performed following the procedure reported in Supplementary Text 10.

## ACKNOWLEDGMENTS

We would like to thank our investors and the executive leadership team of Novel Bio for supporting this work, and Twist Bioscience for the provision of a panel of test plasmids from their catalog of off-the-shelf backbones.

## AUTHOR CONTRIBUTIONS

All authors contributed to design and execution of experiments and analysis of data. MW wrote the paper with support from EW, ML, CO, and NS. MW supervised the research.

