## Supplementary Materials for "Outpacing *E. coli*: Development of *Vibrio natriegens* as a Next-Generation Cloning Host"

### Summary of Contents

**Supplementary Text 1: BLAST of *E. coli recA* against *V. natriegens* genome**

**Supplementary Text 2: Antibiotic Mean Inhibitory Concentrations (MIC) for *V. natriegens***

**Supplementary Text 3: Putative efflux pumps in *V. natriegens* and the impact of their deletions on Kanamycin MIC**

**Supplementary Text 4: Genes identified via differential expression analysis of *V. natriegens* strains grown in the presence or absence of sub-lethal concentrations of Kanamycin**

**Supplementary Text 5: Deletion of capsular polysaccharide, exopolysaccharide, and cell wall-modifying enzyme gene/gene clusters and the impact on Kan MIC**

**Supplementary Text 6: Competent cell stability and reproducibility of batches**

**Supplementary Text 7: Impact of different buffering agents in rich media on plasmid yield**

**Supplementary Text 8: Response Surface Design for Rich Media (liquid) Optimization**

**Supplementary Text 9: Response Surface Design for Rich Media (agar) Optimization**

**Supplementary Text 10: Cloning using Ligation Independent Cloning (LIC)**

**Supplementary Text 11: Plasmids used in this study**

**Supplementary Text 12: Sequences of pCYC series plasmids**

**References**

#### Supplementary Text 1: BLAST of *E. coli recA* against *V. natriegens* genome

The following *E. coli* K12 *recA* sequence was used in a BLAST search:

```
>Ecoli_recA
MAIDENKQKALAAALGQIEKQFGKGSIMRLGEDRSMDVETISTGSLSLDIALGAGGLPMGRIVE
IYGPESSGKTTLTLQVIAAAQREGKTCAFIDAEHALDPIYARKLGVDIDNLLCSQPDTGEQALE
ICDALARSGAVDVIVVDSVAALTPKAEIEGEIGDSHMGLAARMMSQAMRKLAGNLKQSNTLLIF
INQIRMKIGVMFGNPETTTGGNALKFYASVRLDIRRIGAVKEGENVVGSETRVKVVKNKIAAPF
KQAEFQILYGEGINFYGELVDLGVKEKLIEKAGAWYSYKGEKIGQGKANATAWLKDNPETAKEI
EKKVRELLLSNPNSTPDFSVDDSEGVAETNEDF
```

This protein sequence was used as the query in a BLAST against a database containing the DNA sequences of the two *Vibrio natriegens* type strain chromosomes (accession numbers CP016345 and CP016346) using the tblastn algorithm. The following two hits against chromosome I (CP016345) are returned (no hits against chromosome II are observed):

**BLAST results:**

| Accession | Hit start | Hit end | Bit-Score | E Value | Grade | % Identical | Query coverage |
|---|---|---|---|---|---|---|---|
| CP016345 | 2,673,880 | 2,672,897 | 553.132 | 1.43e-180 | 88.7% | 84.5% | 92.92% |
| CP016345 | 2,123,769 | 2,124,032 | 35.809 | 6.87e-03 | 12.7% | 26.7% | 25.50% |

Of the two hits, the first (with an E Value of 1.43e-180) is a strong match for a potential *recA* homolog. The sequence is as follows:

```
>CP016345_recA_homolog
MDENKQKALAAALGQIEKQFGKGSIMRLGDNRAMDVETISTGSLSLDIALGAGGLPMGRI
VEVYGPESSGKTTLTLELIAAAQREGKTCAFIDAEHALDPVYAKKLGVDIDALLVSQPDT
GEQALEICDALARSGAIDVMVVDSVAALTPKAEIEGEMGDSHMGLQARMLSQAMRKLTGN
LKQSNCMCIFINQIRMKIGVMFGNPETTTGGNALKFYASVRLDIRRTGAIKEGDEVVGNE
TRIKVVKNKIAAPFKEANTQIMYGQGFNREGELVDLGVKHKLVEKAGAWYSYNGDKIGQG
KANACNYLREHTEVAQTIDKKLREMLLSPAVAEGPEAGEMPEKKEEEF
```

This is the gene sequence that was pursued for deletion.

**Supplementary Text 2: Antibiotic Mean Inhibitory Concentrations (MIC) for *V. natriegens***

| Antibiotic | MIC (mg/L) |
|---|---|
| Carbenicillin | 1.56 |
| Chloramphenicol | 1.06 |
| Kanamycin | 250 |
| Tetracycline | 3.75 |

The MIC experiment was performed in NBx Cyclone™ growth media in 96-well deep-well plate format, with 1 mL of media supplemented with different concentrations of antibiotics per well. The range of antibiotic concentrations tested were:

- **Carbenicillin** at 100, 50, 25, 12.5, 6.25, 3.13, 1.56, 0.78, 0.39 mg/L
- **Chloramphenicol** at 68, 34, 17, 8.5, 4.5, 2.13, 1.06, 0.53, 0.27, 0.13 mg/L
- **Kanamycin** at 2000, 1000, 500, 250, 125, 62.5, 31.25, 15.63, 7.81, 3.91 mg/L
- **Tetracycline** at 30, 15, 7.5, 3.75, 1.88, 0.94, 0.47, 0.23, 0.12, 0.06 mg/L.

Briefly, each well in columns 2-12 of the 96-well deep-well plate was filled with 1 mL of NBx Cyclone™ growth media containing no antibiotics. Wells A1 and B1 were filled with 2 mL Cyclone™ growth media containing 100 mg/L carbenicillin. Wells C1 and D1 were filled with 2 mL Cyclone™ growth media containing 68 mg/L chloramphenicol. Wells E1 and F1 were filled with 2 mL Cyclone™ growth media containing 2000 mg/L kanamycin. Wells G1 and H2 were filled with 2 mL Cyclone™ growth media containing 30 mg/L tetracycline. A series of 2-fold serial dilution was carried out by transferring 1 mL from each well in column 1 to column 2, mixing, then 1 mL from column 2 to column 3, and so on up to column 10. After that, 1 mL of media was discarded from wells in column 10.

Seed cultures of NBx Cyclone™ were prepared by inoculating cells scrapped from a glycerol stock into 25 mL of growth media without antibiotics and incubating at 37°C with 250 rpm agitation for 8 hours. The culture was then diluted 20-fold with fresh growth media and 20 µL was used to inoculate each well in columns 1to 11 in the MIC experiment deep-well plate described in previous paragraph. Wells in column 11 served as positive control since media in that column did not contain any antibiotics. Wells in column 12 served as negative control since these wells were not inoculated. The MIC experiment deep-well plate was then incubated for 16 hours at 37°C with 900 rpm agitation. After overnight incubation, 100 µL of the cultures in each well in the MIC experiment deep-well plate were transferred to a clear, flat-bottom 96-well plate and cell densities (OD600) were measured by using a microplate reader. The MIC for each antibiotic is the lowest antibiotic concentration in a series that resulted in cell densities no more than 2-fold to average OD600 value of the negative control wells in column 12.

**Supplementary Text 3: Putative efflux pumps in *V. natriegens* and the impact of their deletions on Kanamycin MIC**

We bioinformatically identified putative efflux pumps by first uploading the sequences of each chromosome to the RAST server for annotation[1], and then using the associated SEED Viewer[2] to search for efflux pumps. "Multidrug Resistance Efflux Pumps" were organized under "Resistance to antibiotics and toxic compounds" in the Subsystem Feature Counts menu. The table below shows the locations of each putative efflux pump in the *V. natriegens* genome along with the impact to Kanamycin MIC of the associated knockout strain.

MIC for this and the subsequent datasets was tested by measuring growth at a starting OD600 of 0.01 from an overnight culture, and then grown for 16 hours in a 96 deep-well plate (37°C, 800 RPM) in 500 µL of media (10 g/L tryptone, 5 g/L yeast extract, 633.12 mM NaCl, 9.4 mM KCl, 23 mM MgSO4, 29 mM MgCl2, 9.4 mM CaCl2) at concentrations of Kanamycin 200 to 450 mg/L in 50 mg/L increments. MIC cutoff was established at the Kanamycin concentration where the endpoint OD600 was measured to be <10% of the maximum uninhibited OD600. The MIC was measured for each knockout strain and then compared to the original host strain with MIC Kan of 350 mg/L in this media. The knockout strains for Efflux Pump 3 were not produced in time for the execution of the experiment.

| Efflux Pump | Chromosome | Minimum* | Maximum* | Length | Direction | MIC ratio† |
|---|---|---|---|---|---|---|
| 1 | 1 | 37,476 | 42,315 | 4840 | reverse | 1 |
| 2 | 1 | 419,578 | 420,900 | 1323 | reverse | 1 |
| 3 | 1 | 971,922 | 976,129 | 4208 | forward | 0.778 |
| 4 | 1 | 116,7096 | 1,171,396 | 4301 | forward | N/A |
| 5 | 1 | 1,247,009 | 1,251,192 | 4184 | forward | 1 |
| 6 | 1 | 1,608,125 | 1,609,495 | 1371 | reverse | 1 |
| 7 | 1 | 2,426,342 | 2,427,928 | 1587 | reverse | 1 |
| 8 | 1 | 2,547,184 | 2,548,530 | 1347 | reverse | 0.875 |
| 9 | 1 | 2,572,942 | 2,577,152 | 4211 | reverse | 0.875 |
| 10 | 1 | 2,989,332 | 2,990,717 | 1386 | forward | 0.875 |
| 11 | 2 | 231,737 | 236,097 | 4361 | forward | 1 |
| 12 | 2 | 342,509 | 347,660 | 5152 | reverse | 0.778 |
| 13 | 2 | 1,265,901 | 1,267,241 | 1341 | forward | 1.17 |
| 14 | 2 | 1,386,379 | 1,390,892 | 4514 | forward | 1 |
| 15 | 2 | 1,454,367 | 1,458,695 | 4329 | forward | 0.778 |
| 16 | 2 | 1,468,462 | 1,473,011 | 4550 | reverse | 1 |
| 17 | 2 | 1,587,717 | 1,589,054 | 1338 | reverse | 1 |
| 18 | 2 | 1,622,421 | 1,626,709 | 4289 | forward | 1 |

| | | | | | | |
|---|---|---|---|---|---|---|
| 19 | 2 | 1,922,347 | 1,927,112 | 4766 | reverse | 1.40 |

* Min/Max positions for genes are relative to the sequence numbering used in accession numbers CP016345 and CP016346

† The MIC ratio is a ratio of the Kan MIC (in mg/L) of the control strain to the KO strain. Values close to 1 indicate no change in MIC between the control and the KO strain. Values <1 indicate the MIC of the KO strain is higher (less sensitive to Kanamycin) than the control strain. Values >1 indicate the MIC of the KO strain is lower (more sensitive to Kanamycin) than the control strain.

**Supplementary Text 4: Genes identified via differential expression analysis of *V. natriegens* strains grown in the presence or absence of sub-lethal concentrations of Kanamycin**

The following dataset runs from the Sequence Read Archive were downloaded:

| Dataset run from SRA | Condition* | Replicate |
|---|---|---|
| SRR21618288 | M9Na media + glucose + Kanamycin | 1 |
| SRR21618287 | M9Na media + glucose + Kanamycin | 2 |
| SRR21618292 | M9Na media + glucose | 1 |
| SRR21618291 | M9Na media + glucose | 2 |

*All cultivations were grown in mid-log phase in "M9Na" media, which is standard M9 media supplemented with additional NaCl to more optimally support growth of *V. natriegens*

The above RNA-seq reads were mapped to the *V. natriegens* genome (accession numbers CP016345 and CP016346) and expression levels were calculated in Geneious Prime. Expression levels between the cultivation conditions with and without Kanamycin were compared within Geneious Prime using DESeq2. Data was exported as a CSV file and was sorted by genes that had the highest differential expression between the conditions with and without Kanamycin (log2 ratio). The top 12 genes from this list were knocked out and the impact on Kanamycin MIC relative to the control strain was assessed.

The MIC of KO strains was evaluated relative to the control strain following the procedure outlined in Supplementary Text 3. The knockout strain for aspartate ammonia-lyase was not produced in time for the execution of the experiment.

| Gene annotation | Chromosome | Gene start* | Direction | Differential expression (log2 ratio) | MIC ratio† |
|---|---|---|---|---|---|
| outer membrane protein OmpW | 2 | 1291114 | forward | 8.956 | 1 |
| DUF3149 domain-containing protein | 1 | 788742 | Reverse | 7.971 | 1.17 |
| fusA homolog | 2 | 1446254 | Reverse | 7.359 | 0.875 |
| fumarate reductase | 1 | 2978151 | Forward | 7.045 | 1 |
| transcriptional regulator | 1 | 2024497 | Forward | 6.911 | 1.17 |
| phosphoenolpyruvate carboxykinase (ATP) | 1 | 135378 | Reverse | 6.521 | 1 |
| fumarate reductase (quinol) flavoprotein subunit | 1 | 2975208 | Forward | 6.395 | 0.875 |
| heat-shock protein Hsp20 | 1 | 6622 | Forward | 6.153 | 1 |
| formate dehydrogenase subunit alpha | 2 | 778337 | Reverse | 6.045 | 1.40 |

| | | | | | |
|---|---|---|---|---|---|
| aspartate ammonia-lyase | 1 | 2997536 | Forward | 6.004 | N/A |
| hypothetical protein | 1 | 763645 | Forward | 5.997 | 0.875 |
| universal stress global response regulator UspA | 2 | 792687 | Forward | 5.967 | 0.875 |

* Gene start is relative to the sequence numbering used in accession numbers CP016345 and CP016346

† The MIC ratio is a ratio of the Kan MIC (in mg/L) of the control strain to the KO strain. Values close to 1 indicate no change in MIC between the control and the KO strain. Values <1 indicate the MIC of the KO strain is higher (less sensitive to Kanamycin) than the control strain. Values >1 indicate the MIC of the KO strain is lower (more sensitive to Kanamycin) than the control strain.

Based on an initial literature search, some of the genes in the list were anticipated to have impacts on Kanamycin sensitivity. For example:

- The outer membrane protein OmpW has been shown to participate in *E. coli* with small multidrug resistance proteins in compound efflux[3].
- The *fusA* gene (encoding elongation factor G) is commonly mutated in kanamycin-resistant *E. coli* strains to prevent kanamycin from interfering with protein synthesis[4].
- Amplification of the fumarate reductase operon increased the frequence of persister *E. coli* cells that could survive in the presence of lethal concentrations of antibiotics[5].

However, as the table above shows, none of the 12 knockout strains showed any significant impact on Kanamycin MIC relative to the control strain.

**Supplementary Text 5: Deletion of capsular polysaccharide, exopolysaccharide, and cell wall-modifying enzyme gene/gene clusters and the impact on Kan MIC**

The following genes or operons related to capsule synthesis, exopolysaccharide production, or cell wall-modifying enzymes were deleted in *V. natriegens*, and their impacts on Kanamycin sensitivity assessed.

The MIC of KO strains was evaluated relative to the control strain following the procedure outlined in Supplementary Text 3.

| Gene/gene cluster | Chromosome | Location* | MIC ratio† |
|---|---|---|---|
| *cps* | 2 | 904,848 – 919,729 | 1.40 |
| *syp* | 1 | 1,907,595 – 1,928,982 | 1 |
| *mltG* | 1 | 2,143,632 – 2,144,638 | 1.40 |
| *wbFF* | 1 | 246,835 – 248,568 | 1 |

* Location is relative to the sequence numbering used in accession numbers CP016345 and CP016346

† The MIC ratio is a ratio of the Kan MIC (in mg/L) of the control strain to the KO strain. Values close to 1 indicate no change in MIC between the control and the KO strain. Values <1 indicate the MIC of the KO strain is higher (less sensitive to Kanamycin) than the control strain. Values >1 indicate the MIC of the KO strain is lower (more sensitive to Kanamycin) than the control strain.

**Supplementary Text 6: Competent cell stability and reproducibility of batches**

Commercial chemically competent cells of NBx Cyclone™ were manufactured by Novel Bio. Manufacturing batches show an average transformation efficiency with pUC19 of 4.05E+09 CFU/μg of plasmid DNA (n=7 batches).

A single lot of competent cells (prepared on 06/02/2025) was reserved for long-term stability studies. At regular intervals, aliquots of competent cells from this batch were retrieved and tested in triplicate by transforming with pUC19:

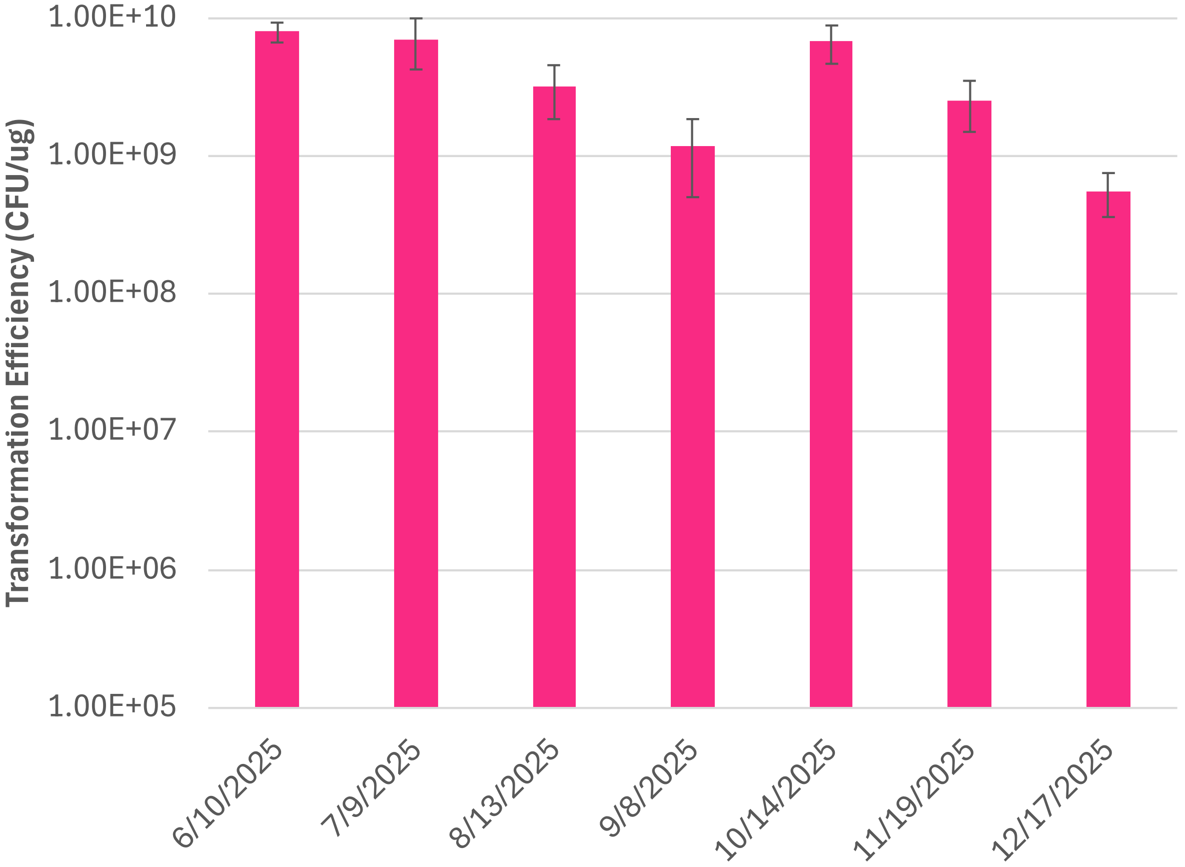

**Supplementary Text 7: Impact of different buffering agents in rich media on plasmid yield**

To evaluate the impact of adding a buffering agent to rich media, media (20 g/L yeast extract, 5 g/L tryptone, 41 mM $MgSO_4$, 428 mM NaCl, 21 mM potassium phosphate) was prepared with either no buffering agent or 150 mM of MES, MOPS, PIPES, or TRIS buffer. All medias were pH adjusted to ~6.4. Colonies of NBx Cyclone™ transformed with pcDNA3.1 were picked into 2 mL of each growth media in wells of a 24-well deep-well plate. The plate was incubated at 37°C with shaking at 300 RPM for 6 hours. Each media was tested in triplicate. 0.3 mL of each culture was harvested and processed through a Qiagen miniprep column as described in the materials and methods section. Plasmid yields were determined using the Qubit fluorometer.

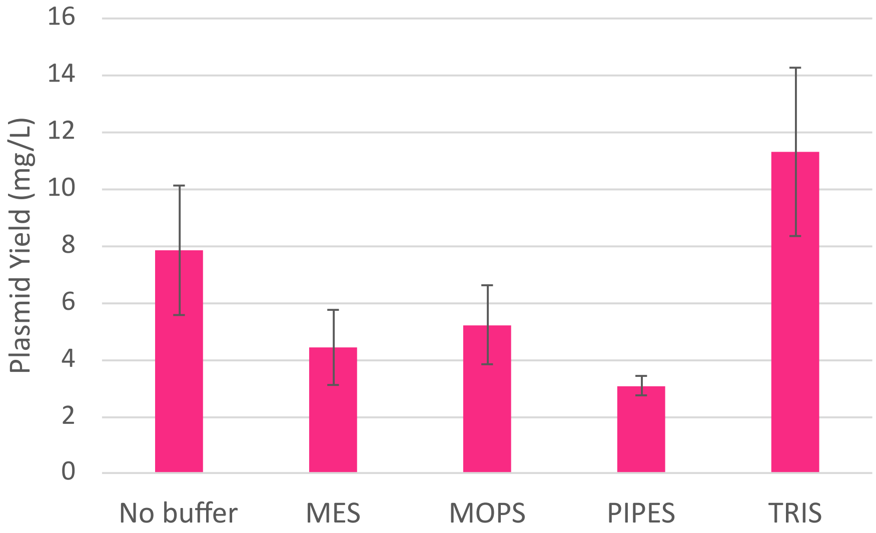

**Supplementary Text 8: Response Surface Design for Rich Media (liquid) Optimization**

To develop an optimized rich media formulation for plasmid DNA production in *V. natriegens*, we employed a response surface methodology using a central composite design. Media components were selected based on the nutritional and physiological requirements of this marine bacterium: yeast extract and vegetable tryptone as complex nitrogen sources; $MgSO_4$, NaCl, and $K_2HPO_4$ to provide essential marine ions and phosphate for nucleotide synthesis; and Tris buffer to maintain pH stability. Component concentration ranges were established using existing *V. natriegens* formulations (e.g., LBv2[6]) and common *E. coli* rich media (LB, TB) as reference points.

The DOE employed the following concentration ranges (g/L): yeast extract (15-30) to ensure sufficient complex nutrients; tryptone (2.5-7.5) with a lower range to determine necessity in the presence of high yeast extract; $MgSO_4$ (4-8) to balance magnesium requirements against precipitation risk with phosphate; NaCl (7.5-22.5, approximately 0.75-2.25%) centered around the 1% optimum observed in preliminary growth studies; and $K_2HPO_4$ (1-3) to evaluate supplementation benefits beyond phosphate provided by yeast extract. This design enabled systematic evaluation of main effects, interactions, and quadratic relationships between components and plasmid yield.

**Response surface design:**

| Condition | Pattern | Yeast Extract (g/L) | Tryptone (g/L) | MgSO4 (g/L) | NaCl (g/L) | K2HPO4 (g/L) | Tris (mM) |
|---|---|---|---|---|---|---|---|
| 1 | ++++− | 30 | 7.5 | 8 | 22.5 | 1 | 150 |
| 2 | 0000A | 22.5 | 5 | 6 | 15 | 4 | 150 |
| 3 | 000a0 | 22.5 | 5 | 6 | 0 | 2 | 150 |
| 4 | −−++− | 15 | 2.5 | 8 | 22.5 | 1 | 150 |
| 5 | A0000 | 37.5 | 5 | 6 | 15 | 2 | 150 |
| 6 | +−−−+ | 30 | 2.5 | 4 | 7.5 | 3 | 150 |
| 7 | −++−− | 15 | 7.5 | 8 | 7.5 | 1 | 150 |
| 8 | −−−−− | 15 | 2.5 | 4 | 7.5 | 1 | 150 |
| 9 | ++−++ | 30 | 7.5 | 4 | 22.5 | 3 | 150 |
| 10 | +−+−− | 30 | 2.5 | 8 | 7.5 | 1 | 150 |
| 11 | 00000 | 22.5 | 5 | 6 | 15 | 2 | 150 |
| 12 | 00000 | 22.5 | 5 | 6 | 15 | 2 | 150 |
| 13 | ++−−− | 30 | 7.5 | 4 | 7.5 | 1 | 150 |
| 14 | +−+++ | 30 | 2.5 | 8 | 22.5 | 3 | 150 |
| 15 | 0000a | 22.5 | 5 | 6 | 15 | 0 | 150 |
| 16 | +−−+− | 30 | 2.5 | 4 | 22.5 | 1 | 150 |
| 17 | 00000 | 22.5 | 5 | 6 | 15 | 2 | 150 |
| 18 | 00000 | 22.5 | 5 | 6 | 15 | 2 | 150 |
| 19 | 0a000 | 22.5 | 0 | 6 | 15 | 2 | 150 |
| 20 | −−+−+ | 15 | 2.5 | 8 | 7.5 | 3 | 150 |
| 21 | −−−++ | 15 | 2.5 | 4 | 22.5 | 3 | 150 |
| 22 | −+−+− | 15 | 7.5 | 4 | 22.5 | 1 | 150 |
| 23 | +++−+ | 30 | 7.5 | 8 | 7.5 | 3 | 150 |
| 24 | 00000 | 22.5 | 5 | 6 | 15 | 2 | 150 |
| 25 | a0000 | 7.5 | 5 | 6 | 15 | 2 | 150 |
| 26 | 000A0 | 22.5 | 5 | 6 | 30 | 2 | 150 |
| 27 | 0A000 | 22.5 | 10 | 6 | 15 | 2 | 150 |
| 28 | 00000 | 22.5 | 5 | 6 | 15 | 2 | 150 |
| 29 | −+−−+ | 15 | 7.5 | 4 | 7.5 | 3 | 150 |
| 30 | 00a00 | 22.5 | 5 | 2 | 15 | 2 | 150 |
| 31 | −++++ | 15 | 7.5 | 8 | 22.5 | 3 | 150 |
| 32 | 00A00 | 22.5 | 5 | 10 | 15 | 2 | 150 |

Each of the 32 conditions were formulated, and 2 mL was added to wells of 24-well deep-well plates (2 plates total; each condition n=1, center point n=6 to assess variability). Carbenicillin was added to a concentration of 50 μg/mL. The wells were inoculated with 5 μL of an overnight culture of NBx Cyclone™ harboring plasmid pcDNA3.1 grown in the media with Tris described in Supplementary Text 6. The cultures were grown for 4 hours at 37°C with shaking at 300 RPM and humidity at 80%. At 4 hours, 0.2 mL of each culture was harvested and processed through a Qiagen miniprep column as described in the materials and methods section. Plasmid yields were determined using the Qubit fluorometer. OD600 was measured spectroscopically on a Nanodrop UV/Vis spectrophotometer using the cuvette setting.

Here are the results sorted by plasmid yield:

**Response surface design with OD600 and plasmid yield data (sorted by yield):**

| Condition | Pattern | Yeast Extract (g/L) | Tryptone (g/L) | MgSO4 (g/L) | NaCl (g/L) | K2HPO4 (g/L) | Tris (mM) | OD600 | Plasmid yield (mg/L) |
|---|---|---|---|---|---|---|---|---|---|
| 4 | −−++− | 15 | 2.5 | 8 | 22.5 | 1 | 150 | 5.6 | 1.32 |
| 21 | −−−++ | 15 | 2.5 | 4 | 22.5 | 3 | 150 | 4.4 | 1.18 |
| 15 | 0000a | 22.5 | 5 | 6 | 15 | 0 | 150 | 6 | 1.12 |
| 25 | a0000 | 7.5 | 5 | 6 | 15 | 2 | 150 | 6 | 1.01 |
| 1 | ++++− | 30 | 7.5 | 8 | 22.5 | 1 | 150 | 3.8 | 0.998 |
| 22 | −+−+− | 15 | 7.5 | 4 | 22.5 | 1 | 150 | 5.6 | 0.996 |
| 16 | +−−+− | 30 | 2.5 | 4 | 22.5 | 1 | 150 | 4.6 | 0.928 |
| 19 | 0a000 | 22.5 | 0 | 6 | 15 | 2 | 150 | 4.6 | 0.926 |
| 24 | 00000 | 22.5 | 5 | 6 | 15 | 2 | 150 | 5.2 | 0.926 |
| 17 | 00000 | 22.5 | 5 | 6 | 15 | 2 | 150 | 5.4 | 0.912 |
| 18 | 00000 | 22.5 | 5 | 6 | 15 | 2 | 150 | 5.2 | 0.91 |
| 11 | 00000 | 22.5 | 5 | 6 | 15 | 2 | 150 | 5.4 | 0.89 |
| 12 | 00000 | 22.5 | 5 | 6 | 15 | 2 | 150 | 5.8 | 0.874 |
| 30 | 00a00 | 22.5 | 5 | 2 | 15 | 2 | 150 | 5 | 0.86 |
| 28 | 00000 | 22.5 | 5 | 6 | 15 | 2 | 150 | 5.6 | 0.854 |
| 31 | −++++ | 15 | 7.5 | 8 | 22.5 | 3 | 150 | 4.2 | 0.81 |
| 7 | −++−− | 15 | 7.5 | 8 | 7.5 | 1 | 150 | 4.8 | 0.794 |
| 9 | ++−++ | 30 | 7.5 | 4 | 22.5 | 3 | 150 | 3.2 | 0.794 |
| 27 | 0A000 | 22.5 | 10 | 6 | 15 | 2 | 150 | 5.4 | 0.788 |
| 32 | 00A00 | 22.5 | 5 | 10 | 15 | 2 | 150 | 4.8 | 0.782 |
| 2 | 0000A | 22.5 | 5 | 6 | 15 | 4 | 150 | 4.4 | 0.774 |
| 14 | +−+++ | 30 | 2.5 | 8 | 22.5 | 3 | 150 | 3.2 | 0.76 |
| 20 | −−+−+ | 15 | 2.5 | 8 | 7.5 | 3 | 150 | 3.6 | 0.754 |
| 26 | 000A0 | 22.5 | 5 | 6 | 30 | 2 | 150 | 3.4 | 0.734 |
| 5 | A0000 | 37.5 | 5 | 6 | 15 | 2 | 150 | 3.8 | 0.714 |
| 10 | +−+−− | 30 | 2.5 | 8 | 7.5 | 1 | 150 | 4.6 | 0.656 |
| 13 | ++−−− | 30 | 7.5 | 4 | 7.5 | 1 | 150 | 4 | 0.614 |
| 6 | +−−−+ | 30 | 2.5 | 4 | 7.5 | 3 | 150 | 3.4 | 0.538 |
| 29 | −+−−+ | 15 | 7.5 | 4 | 7.5 | 3 | 150 | 4.4 | 0.534 |
| 23 | +++−+ | 30 | 7.5 | 8 | 7.5 | 3 | 150 | 3.2 | 0.522 |
| 3 | 000a0 | 22.5 | 5 | 6 | 0 | 2 | 150 | 0.6 | 0 |
| 8 | −−−−− | 15 | 2.5 | 4 | 7.5 | 1 | 150 | 1.6 | 0 |

A strong statistically significant model was created with an $R^2 = 0.96$, RMSE of 0.078 mg/L, and P-value <0.0001. Among the 14 significant model terms, NaCl concentration exhibited the strongest linear effect, followed by a significant quadratic term for NaCl, indicating an optimal concentration range. The $MgSO_4$ x $K_2HPO_4$ interaction was the third most influential factor, suggesting synergistic or antagonistic effects between these salt components on plasmid yield. The quadratic term for $K_2HPO_4$ was excluded due to low P value (0.2). Counterintuitively, both yeast extract and tryptone showed negative linear effects on plasmid yield at 4 hours, with optima at 7.5 g/L and 0 g/L, respectively. This indicates that moderate nutrient limitation enhances early plasmid accumulation in this fast-growing marine bacterium, possibly by reducing acetate or other metabolic byproducts that accumulate during rapid growth on rich media, or by shifting the balance between cell division and plasmid replication.

**Analysis of model in JMP, and predicted optimum condition:**

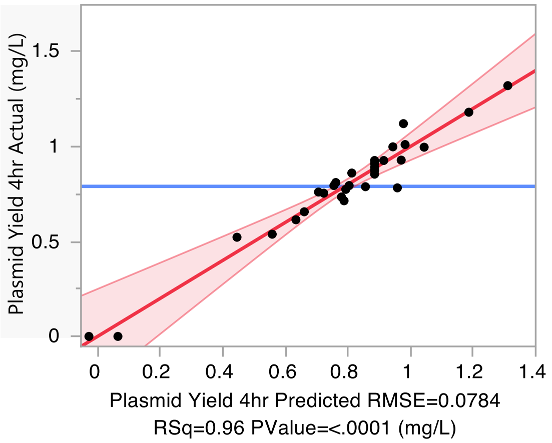


| Source | Logworth | | PValue |
|---|---|---|---|
| NaCl(7.5,22.5) | 8.997 | | 0.00000 |
| NaCl*NaCl | 6.864 | | 0.00000 |
| MgSO4*K2HPO4 | 3.492 | | 0.00032 |
| Tryptone*NaCl | 2.552 | | 0.00281 |
| Yeast Extract*NaCl | 2.458 | | 0.00348 |
| Tryptone*K2HPO4 | 2.447 | | 0.00358 |
| MgSO4*NaCl | 2.388 | | 0.00409 |
| NaCl*K2HPO4 | 2.190 | | 0.00645 |
| Yeast Extract(15,30) | 2.113 | | 0.00771 |
| Yeast Extract*MgSO4 | 1.971 | | 0.01070 |
| K2HPO4(1,3) | 1.962 | | 0.01091 |
| Yeast Extract*K2HPO4 | 1.531 | | 0.02945 |
| MgSO4(4,8) | 1.431 | | 0.03703 |
| Tryptone*MgSO4 | 1.282 | | 0.05218 |
| Tryptone(2.5,7.5) | 0.425 | | 0.37582 |
| K2HPO4*K2HPO4&Excluded | . | | . |

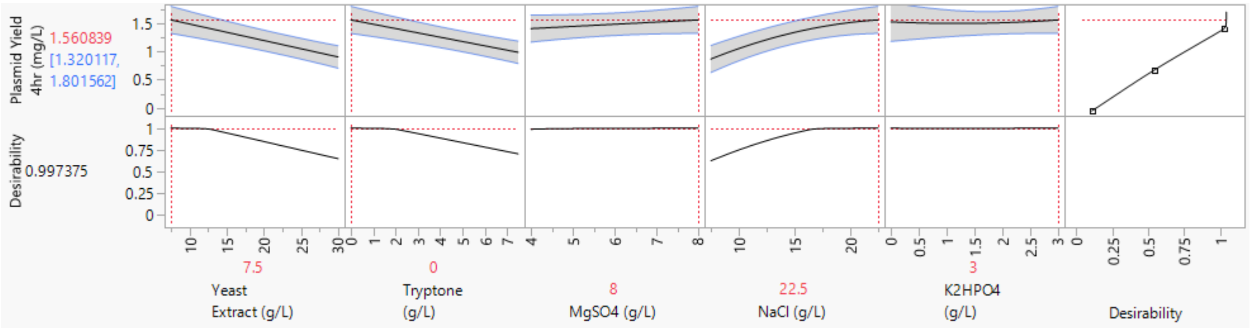


To further validate the model, the top three performers (conditions 4, 15, and 21) along with a low-performing control (condition 26) were retested to confirm reproducibility of the performance ranking.

For this second experiment, NBx Cyclone™ harboring either pUC19 or pACYC184 was used as the strain. Strains were streaked out on agar plates containing antibiotic (Carbenicillin for pUC19 and Chloramphenicol for pACYC184) and incubated overnight at 37°C to generate colonies. Single colonies were picked into 4 mL cultures in wells of 24-well deep-well blocks of the respective media (two cultures of each media were each inoculated from an independent colony). The cultures were incubated at 37°C with 80% humidity and agitation at 300 RPM. Samples were taken for OD600 and yield determination at 4, 6, and 8 hours. Plasmid yields were determined using the Qubit fluorometer, and were used to calculate volumetric yield (mg/L). OD600 was measured spectroscopically on a Nanodrop UV/Vis spectrophotometer using the cuvette setting. Specific yield was calculated by dividing yield (mg/L) by OD600 (giving units of mg/L/OD).

**Results for pUC19:**

| Plasmid | Condition | Time (hrs) | OD600 | OD (st dev) | Yield (mg/L) | Yield (st dev) | Spec Yld (mg/L/OD) | Spec Yld (st dev) |
|---|---|---|---|---|---|---|---|---|
| pUC19 | 4 | 4 | 6.35 | 0.35355339 | 0.62 | 0.87681241 | 0.1 | 0.141421356 |
| pUC19 | 15 | 4 | 6.2 | 0.84852814 | 0 | 0 | 0 | 0 |
| pUC19 | 21 | 4 | 6.45 | 0.35355339 | 0 | 0 | 0 | 0 |
| pUC19 | 26 | 4 | 5.25 | 0.07071068 | 0 | 0 | 0 | 0 |
| pUC19 | 4 | 6 | 9.65 | 0.07071068 | 4.425 | 0.09192388 | 0.455 | 0.007071068 |
| pUC19 | 15 | 6 | 12.45 | 0.91923882 | 3.195 | 1.03944697 | 0.255 | 0.06363961 |
| pUC19 | 21 | 6 | 10.8 | 0.42426407 | 4.55 | 0.15556349 | 0.42 | 0.028284271 |
| pUC19 | 26 | 6 | 10.25 | 0.21213203 | 2.935 | 0.19091883 | 0.29 | 0.014142136 |
| pUC19 | 4 | 8 | 10.5 | 0.70710678 | 10.575 | 1.59099026 | 1.015 | 0.219203102 |
| pUC19 | 15 | 8 | 15.25 | 1.06066017 | 11 | 1.76776695 | 0.725 | 0.16263456 |
| pUC19 | 21 | 8 | 10.75 | 0.35355339 | 11.625 | 0.31819805 | 1.085 | 0.06363961 |
| pUC19 | 26 | 8 | 13.5 | 0.70710678 | 10.8 | 0.84852814 | 0.805 | 0.106066017 |

**Results for pACYC184:**

| Plasmid | Condition | Time (hrs) | OD600 | OD (st dev) | Yield (mg/L) | Yield (st dev) | Spec Yld (mg/L/OD) | Spec Yld (st dev) |
|---|---|---|---|---|---|---|---|---|
| pACYC184 | 4 | 4 | 3.1 | 3.53553391 | 1.13 | 1.59806133 | 0.2 | 0.282842712 |
| pACYC184 | 15 | 4 | 6.8 | 0.56568542 | 2.495 | 0.54447222 | 0.365 | 0.049497475 |
| pACYC184 | 21 | 4 | 6.65 | 1.20208153 | 3.055 | 0.82731493 | 0.455 | 0.035355339 |
| pACYC184 | 26 | 4 | 2 | 0.28284271 | 0 | 0 | 0 | 0 |
| pACYC184 | 4 | 6 | 9.15 | 4.03050865 | 4.59 | 3.33754401 | 0.465 | 0.16263456 |
| pACYC184 | 15 | 6 | 14.2 | 0.14142136 | 9.5 | 0.91923882 | 0.67 | 0.056568542 |
| pACYC184 | 21 | 6 | 12.55 | 0.77781746 | 6.95 | 0.21213203 | 0.555 | 0.021213203 |
| pACYC184 | 26 | 6 | 8.95 | 0.07071068 | 3.65 | 0.35355339 | 0.405 | 0.035355339 |
| pACYC184 | 4 | 8 | 14.25 | 0.35355339 | 9.725 | 3.85373196 | 0.68 | 0.254558441 |
| pACYC184 | 15 | 8 | 18 | 0.70710678 | 11.65 | 0.70710678 | 0.65 | 0.014142136 |
| pACYC184 | 21 | 8 | 17 | 1.41421356 | 12.55 | 0.07071068 | 0.74 | 0.070710678 |
| pACYC184 | 26 | 8 | 15.27 | 0.38183766 | 9.125 | 1.4495689 | 0.595 | 0.077781746 |

As can be seen from the above datasets, condition 26 reproduced as a low performer at early timepoints, while conditions 4, 15, and 21 reproduced as high performers. Volumetric yields and specific yields are generally higher in condition 21 across timepoints and plasmids. As a result, condition 21 was selected as our best tested media to move forward.

As a final experiment, condition 21 was compared to the predicted optimal media formulation from the DOE (7.5 g/L yeast extract, 8 g/L $MgSO_4$, 22.5 g/L NaCl, 3 g/L $K_2HPO_4$, 150 mM Tris, pH 6.4). NBx CyClone™ harboring pcDNA3.1 was cultivated in 2 mL culture volume in 24-well deep-well plates in both condition 21 as well as the model-predicted media in triplicate (n=3). Samples were processed at 4, 6, and 8 hours as above, and specific yields (mg/L/OD) and volumetric yields (mg/L) were calculated for each:

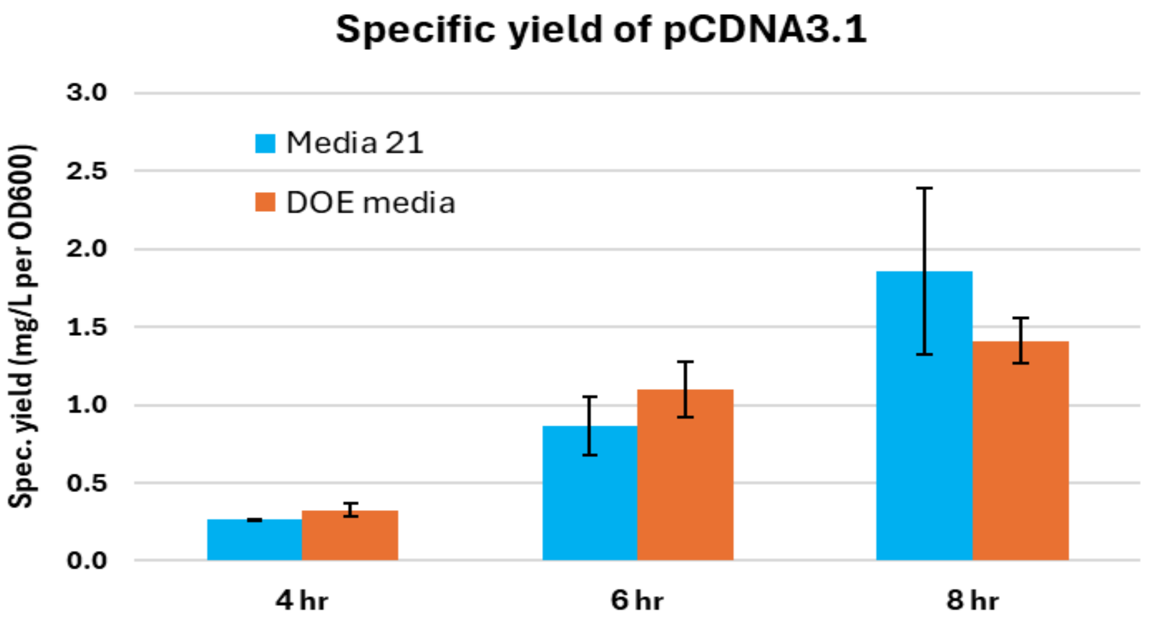


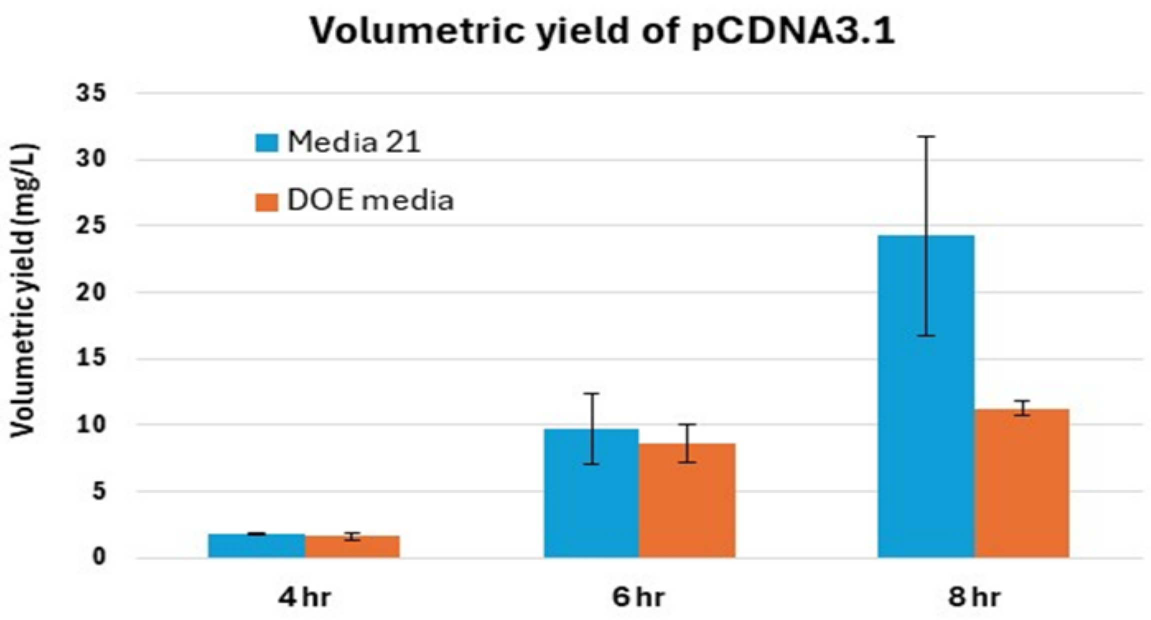


Both condition 21 and the DOE-predicted media formulations performed similarly at 4 and 6 hours. At 8 hours, condition 21 showed an improvement in both metrics, particularly volumetric yield. This is likely due to the inclusion of additional nutrients (tryptone and more yeast extract) in condition 21. As a result, we decided to proceed with selecting condition 21 as our NBx Cyclone™ media.

**Supplementary Text 9: Response Surface Design for Rich Media (agar) Optimization**

Rationale for media components and execution of DOE design followed a similar design philosophy as used for the liquid rich media (see Supplementary Text 8) with the addition of additional salts (KCl and $CaCl_2$). The DOE employed the following concentration ranges: yeast extract (1-9 g/L) and tryptone (1-19 g/L); KCl (0-18.8 mM), $MgSO_4$ (23-46 mM), $CaCl_2$ (0-18.8 mM), $MgCl_2$ (0-58 mM). The midpoints for each concentration were derived from a media composition (LB Tris) that was chosen to be tested as an agar media based on successful preliminary testing as a rich liquid media. 15 g/L agar (1.5%) was used for all agar media configurations, while LB Tris recipe also contained 15.87 mM Potassium Phosphate (pH 6.4) and 113.38 mM Tris (pH 7).

**Response surface design:**

| Condition | Yeast Extract (g/L) | Tryptone (g/L) | NaCl (mM) | KCl (mM) | MgSO4 (mM) | MgCl2 (mM) | CaCl2 (mM) |
|---|---|---|---|---|---|---|---|
| **1** | 1 | 1 | 633.12 | 0 | 46 | 0 | 9.4 |
| **2** | 1 | 19 | 171.12 | 0 | 23 | 58 | 18.8 |
| **3** | 1 | 1 | 171.12 | 18.8 | 0 | 58 | 0 |
| **4** | 1 | 19 | 171.12 | 0 | 46 | 29 | 0 |
| **5** | 1 | 1 | 633.12 | 0 | 0 | 58 | 18.8 |
| **6** | 1 | 1 | 171.12 | 18.8 | 46 | 0 | 18.8 |
| **7** | 1 | 19 | 633.12 | 9.4 | 0 | 0 | 0 |
| **8** | 1 | 10 | 633.12 | 18.8 | 46 | 58 | 0 |
| **9** | 1 | 19 | 405.12 | 18.8 | 0 | 0 | 18.8 |
| **10** | 5 | 19 | 633.12 | 18.8 | 46 | 58 | 18.8 |
| **11** | 5 | 10 | 405.12 | 9.4 | 23 | 29 | 9.4 |
| **12** | 5 | 1 | 171.12 | 0 | 0 | 0 | 0 |
| **13** | 9 | 10 | 171.12 | 0 | 0 | 0 | 18.8 |
| **14** | 9 | 19 | 633.12 | 0 | 46 | 0 | 18.8 |
| **15** | 9 | 19 | 171.12 | 18.8 | 0 | 58 | 9.4 |
| **16** | 9 | 1 | 633.12 | 18.8 | 0 | 29 | 18.8 |
| **17** | 9 | 1 | 633.12 | 18.8 | 23 | 0 | 0 |
| **18** | 9 | 1 | 405.12 | 0 | 46 | 58 | 0 |
| **19** | 9 | 19 | 633.12 | 0 | 0 | 58 | 0 |
| **20** | 9 | 1 | 171.12 | 9.4 | 46 | 58 | 18.8 |
| **21** | 9 | 19 | 171.12 | 18.8 | 46 | 0 | 0 |
| **LB** | 5 | 10 | 171.12 | 0 | 0 | 0 | 0 |

Transformations of competent cells with pUC19 were plated and incubated at 37°C for 16 hours on the respective agar designs. Colony size (scored 1-4, with 4 corresponding to fastest growth) and transformation efficiency (number of colonies from 3 µL of transformation plated) were recorded for each plate:

| Condition | Colonies | Colony Size |
|---|---|---|
| **1** | 450 | 2 |
| **2** | 523 | 4 |
| **3** | 28 | 2 |
| **4** | 87 | 3 |
| **5** | 451 | 2 |
| **6** | 598 | 2 |
| **7** | 50 | 3 |
| **8** | 272 | 3 |
| **9** | 400 | 1 |
| **10** | 780 | 3 |
| **11** | 674 | 4 |
| **12** | 0 | 1 |
| **13** | 0 | 1 |
| **14** | 716 | 3 |
| **15** | 579 | 4 |
| **16** | 730 | 3 |
| **17** | 234 | 3 |
| **18** | 269 | 3 |
| **19** | 483 | 3 |
| **20** | 580 | 4 |
| **21** | 257 | 3 |
| **LB** | 151 | 3 |

Colony size and transformation efficiency were evaluated as response variables in the model, enabling systematic evaluation of main effects, interactions, and quadratic relationships between growth rate (colony size) and transformation efficiency (number of colonies).

**Analysis of model in JMP (colony count):**

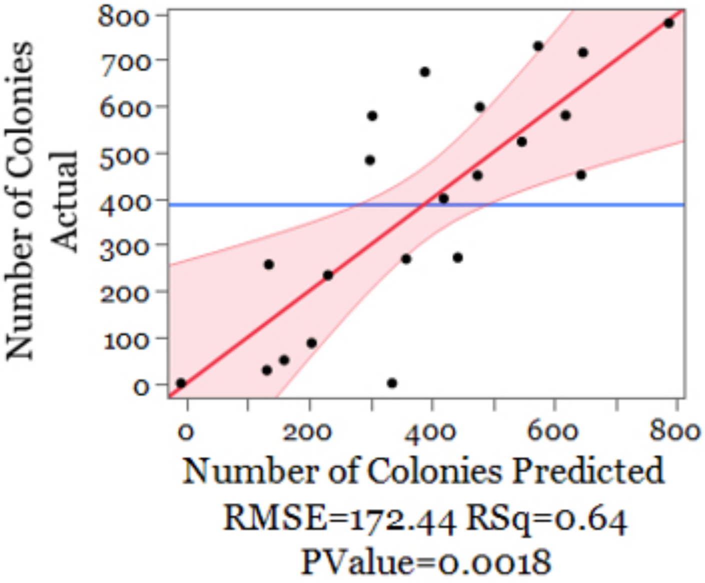


| Term | Estimate | Std Error | t Ratio | Prob>\|t\| | Uncoded Estimate |
|---|---|---|---|---|---|
| Intercept | 388.61905 | 37.62958 | 10.33 | <.0001* | -71.46649 |
| NaCl(171.12,633.12) | 84.111111 | 40.64459 | 2.07 | 0.0551 | 0.3641174 |
| MgSO4(0,46) | 71.555556 | 40.64459 | 1.76 | 0.0974 | 3.1111111 |
| MgCl2(0,58) | 70 | 40.64459 | 1.72 | 0.1043 | 2.4137931 |
| CaCl2(0,18.8) | 172.11111 | 40.64459 | 4.23 | 0.0006* | 18.309693 |

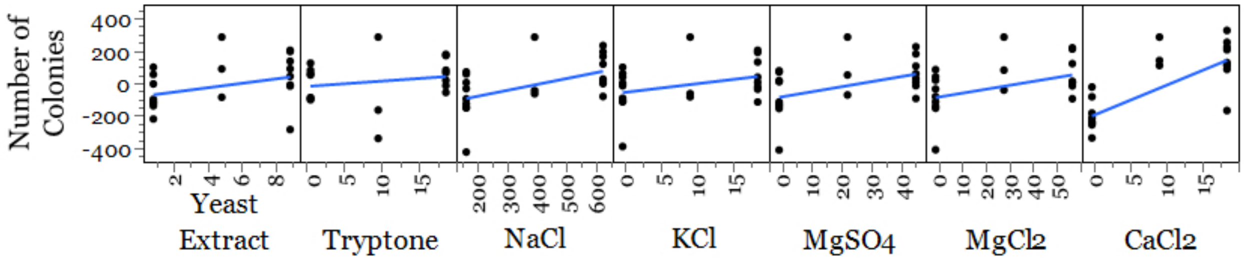


Based on the model, increasing $CaCl_2$ in the agar formulations had the largest influence on improving transformation efficiency, with increasing NaCl, $MgSO_4$, and $MgCl_2$ also having an effect.

**Analysis of model in JMP (colony size):**

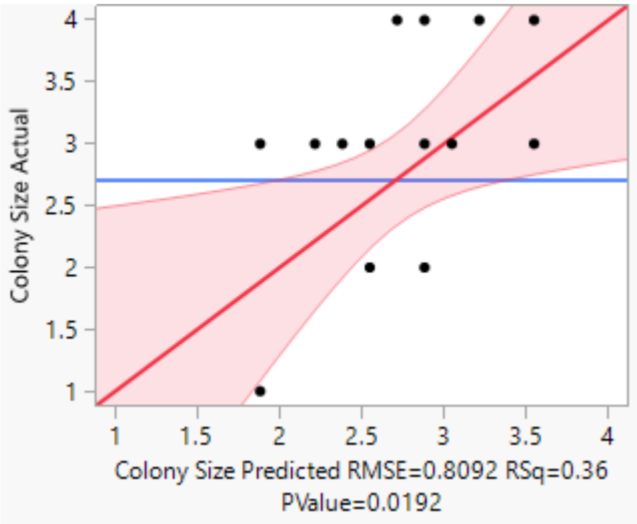


| Term | Estimate | Std Error | t Ratio | Prob>\|t\| |
|---|---|---|---|---|
| Intercept | 1.8809524 | 0.322382 | 5.83 | <.0001* |
| MgSO4 (mM) | 0.0144928 | 0.008292 | 1.75 | 0.0975 |
| MgCl2 (mM) | 0.0172414 | 0.006577 | 2.62 | 0.0173* |

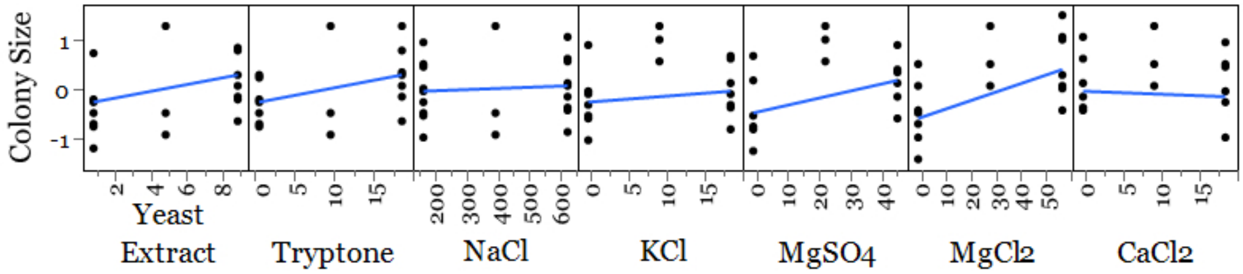


Increasing $MgCl_2$ showed to be the largest influencing factor on increasing colony growth rate, with increasing $MgSO_4$, yeast extract, and tryptone also having an effect.

Follow-up experiments resulted in the final NBx CyClone™ agar formulation, which is composed of 15 g/L agar, 5 g/L yeast extract, 10 g/L vegetable tryptone, 19.54 g/L sodium chloride, 3.82 g/L magnesium sulfate (anhydrous), 117 mM Tris (from a 1 M, pH 7 stock). This formulation gave a good balance between transformation efficiency and colony growth. The exclusion of KCl, $K_2HPO_4$, and $CaCl_2$ from the final formulation eliminated precipitation events during autoclaving the agar media while showing no noticeable decrease in performance

**Supplementary Text 10: Cloning using Ligation Independent Cloning (LIC)**

Primers LIC-vector-F and LIC-vector-R (see table below) were used to amplify the backbone of pCYC02 (plasmid sequence in Supplementary Text 12). Primers LIC-insert-F and LIC-insert-R (see table below) were used to amplify a GFP-expression cassette from an internal plasmid (pCYC03-GFP).

**Primers used for LIC:**

| Primer Name | LIC homology region | Primer region | Full primer (5' to 3') |
|---|---|---|---|
| LIC-insert-F | GAGAGTGAGGTATTGA GGGT | CTATTCAGGGATTGGT AAAAC | GAGAGTGAGGTATTGAGGGTC TATTCAGGGATTGGTAAAAC |
| LIC-insert-R | GAGGTTAGAGGAGAGT TAGAG | CATCCACGCCTGAGAT CTAG | GAGGTTAGAGGAGAGTTAGAG CATCCACGCCTGAGATCTAG |
| LIC-vector-F | ACCCTCAATACCTCAC TCTC | GGCGTAATCATGGTCA TAGC | ACCCTCAATACCTCACTCTCG GCGTAATCATGGTCATAGC |
| LIC-vector-R | CTCTAACTCTCCTCTA ACCTC | GTATATCTGGCCCGTA CAT | CTCTAACTCTCCTCTAACCTC GTATATCTGGCCCGTACAT |

Q5 Hot Start High-Fidelity 2X master mix was used in both PCR reactions. The amplicons from both reactions were gel-purified. The pCYC02 amplicon (0.1 pmole) was mixed 2.5 mM dCTP, 5 mM DTT, 0.5 µL T4 DNA polymerase (NEB) in 1X NEB Buffer 2.1. The GFP amplicon (0.1 pmole) was mixed with 2.5 mM dGTP, 5 mM DTT, 0.5 µL T4 DNA polyemerase in 1X NEB Buffer 2.1. Both of the T4 DNA polymerase reactions were incubated at 23°C for 40 min, followed by 20 min at 75°C. 3 µL of the GFP-T4 DNA polymerase reaction was mixed with 1 µL of the pCYC02-T4 DNA polymerase reaction and incubated at 23°C for 25 min. 2 µL of 50 mM EDTA was then added to the mixture and incubated at 23°C for 5 additional min. 3 µL of this assembly was used to transform NBx CyClone™ chemical competent cells.

To compare LIC assembly to Gibson assembly, the pCYC02 amplicon amplified with LIC-vector-F and -R, and the GFP amplicon amplified with LIC-insert-F and -R were also assembled via Gibson Assembly using the NEBuilder HiFi DNA Assembly protocol (the primer design used in this experiment was intentionally designed to be compatible with both LIC and Gibson Assembly). The assembly product was transformed into NBx CyClone™ chemically competent cells.

Transformations were recovered per the protocol in the Materials & Methods section of the paper, and plated out on agar plates. Various dilutions of the recovery reaction were plated to ensure pickable, well-isolated colonies. 8 colonies from each transformation were randomly picked to inoculate 2-mL of NBx CyClone™ media with chloramphenicol. The cultures were incubated at 37°C at 250 RPM for 5 hours, after which 1 mL of cells were spun down. Agar plates and pellets were visualized under a blue light transilluminator for GFP expression:

**Visualization of GFP expression on agar plates and in cell pellets from cultures.**

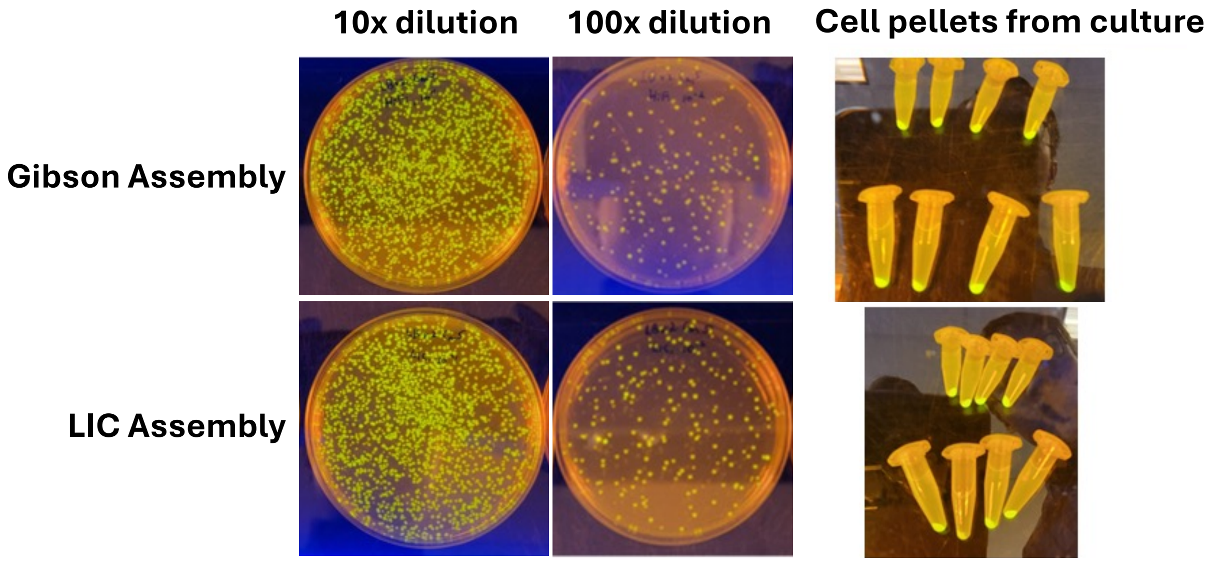


As can be seen from the image above, the majority of colonies from both assembly reactions (as well as all 8 cell pellets from isolated colonies) emitted green fluorescence under blue light, demonstrating that the GFP cassette was successfully cloned into pCYC02 using both approaches. Subsequent sequencing results of plasmids further supported this conclusion.

**Supplementary Text 11: Plasmids used in this study**

The following table contains more details on the plasmids used in this study:

| Plasmid name | Size (bp) | Antibiotic | Origin of replication | Source | Application |
|---|---|---|---|---|---|
| pUC19 | 2,686 | Amp | pUC ori | Internal | General cloning |
| pcDNA3.1 | 5,428 | Amp | pUC† | Thermo | Mammalian protein expression |
| pACYC184 | 4,245 | Chlor | p15a | Internal | General cloning |
| gWIZ-GFP | 5,877 | Kan | pUC | Fisher Scientific | Mammalian protein expression |
| pVAX | 2,999 | Kan | pUC† | Thermo | DNA vaccines |
| pET28a with enzyme (BxbI) CDS | 6,748 | Kan | pBR322 (low copy) | Internal | Bacterial protein expression |
| pTwist Chlor Medium Copy | 2,664 | Chlor | p15a | Twist Bioscience | General cloning |
| pTwist Chlor High Copy | 2,724 | Chlor | pMB1 (pUC) | Twist Bioscience | General cloning |
| pET blank (KAN)* | 5,691 | Kan | pBR322 (low copy) | Twist Bioscience | Bacterial protein expression |
| pET blank (AMP)* | 6,057 | Amp | pBR322 (low copy) | Twist Bioscience | Bacterial protein expression |
| pTwist CMV BG WPRE Neo* | 7,456 | Amp | pUC | Twist Bioscience | Mammalian protein expression |
| pTwist CMV OriP* | 5,631 | Amp | pUC | Twist Bioscience | Mammalian protein expression |
| pTwist Lenti CAG Puro* | 9,832 | Amp | pUC | Twist Bioscience | Stable gene integration via lentivirus |
| pTwist Lenti TRE Puro* | 8,384 | Amp | pUC | Twist Bioscience | Stable gene integration via lentivirus |
| pTwist IVT hBG 50A* | 3,259 | Kan | pUC | Twist Bioscience | In vitro transcription for mRNA production |
| pTwist cDNA BG* | 7,236 | Amp | pUC | Twist Bioscience | Mammalian protein expression |

*All plasmids from Twist contained the open reading frame for Red Fluorescent Protein (RFP) as an insert sequence
†The origin is highly homologous to pUC19, differing only in a few positions

**Supplementary Text 12: Sequences of pCYC series plasmids**

Plasmid sequences for pCYC01 – AmpR, pCYC02 – CmR, and pCYC03 – KanR provided in FASTA format:

```
>pCYC01_AmpR
TTGAGATCCTTTTTTTCTGCGCGTAATCTGCTGCTTGCAAACAAAAAAACCACCGCTACCAGCG
GTGGTTTGTTTGCCGGATCAAGAGCTACCAACTCTTTTTCCGAAGGTAACTGGCTTCAGCAGAG
CGCAGATACCAAATACTGTTCTTCTAGTGTAGCCGTAGTTAGGCCACCACTTCAAGAACTCTGT
AGCACCGCCTACATACCTCGCTCTGCTAATCCTGTTACCAGTGGCTGCTGCCAGTGGCGATAAG
TCGTGTCTTACCGGGTTGGACTCAAGACGATAGTTACCGGATAAGGCGCAGCGGTCGGGCTGAA
CGGGGGGTTCGTGCACACAGCCCAGCTTGGAGCGAACGACCTACACCGAACTGAGATACCTACA
GCGTGAGCTATGAGAAAGCGCCACGCTTCCCGAAGGGAGAAAGGCGGACAGGTATCCGGTAAGC
GGCAGGGTCGGAACAGGAGAGCGCACGAGGGAGCTTCCAGGGGGAAACGCCTGGTATCTTTATA
GTCCTGTCGGGTTTCGCCACCTCTGACTTGAGCGTCGATTTTTGTGATGCTCGTCAGGGGGGCG
GAGCCTATGGAAAAACGCCAGCAACGCGGCCTTTTTACGGTTCCTGGCCTTTTGCTGGCCTTTT
GCTCACATGTTCTTTCCTGCGTTATCCCCTGATTCTGTGGATAACCGTATTACCGCCTTTGAGT
GAGCTGATACCGCTCGCCGCAGCCGAACGACCGAGCGCAGCGAGTCAGTGAGCGAGGAAGCGGA
AGAGCGCCCAATACGCAAACCGCCTCTCCCCGCGCGTTGGCCGATTCATTAATGCAGCTGGCAC
GACAGGTTTCCCGACTGGAAAGCGGGCAGTGAGCGCAACGCAATTAATGTGAGTTAGCTCACTC
ATTAGGCACCCCAGGCTTTACACTTTATGCTTCCGGCTCGTATGTTGTGTGGAATTGTGAGCGG
ATAACAATTTCACACAGGAAACAGCTATGACCATGATTACGCCAAGCTCCTGGAGACCAGCATG
GCTTGCAGTGGCGTTGGTCTCCGGCAACGCGTATATCTGGCCCGTACATCGCGAAGCAGCGCAA
AACGCCTAACCCTAAGCAGATTCTTCATGCAATTGTCGGTCAAGCCTTGCCTTGTTGTAGCTTA
AATTTTGCTCGCGCACTACTCAGCGACCTCCAACACACAAGCAGGGAGCAGATACTGGCTTAAC
TATGCGGCATCAGAGCAGATTGTACTGAGAGTGCACCATAGGGGATCGGGAGATCTCCCGATCC
GTCGACGTCAGGTGGCACTTTTCGGGGAAATGTGCGCGGAACCCCTATTTGTTTATTTTTCTAA
ATACATTCAAATATGTATCCGCTCATGAGACAATAACCCTGATAAATGCTTCAATAATATTGAA
AAAGGAAGAGTATGAGTATTCAACATTTCCGTGTCGCCCTTATTCCCTTTTTTGCGGCATTTTG
CCTTCCTGTTTTTGCTCACCCAGAAACGCTGGTGAAAGTAAAAGATGCTGAAGATCAGTTGGGT
GCACGAGTGGGTTACATCGAACTGGATCTCAACAGCGGTAAGATCCTTGAGAGTTTTCGCCCCG
AAGAACGTTTTCCAATGATGAGCACTTTTAAAGTTCTGCTATGTGGCGCGGTATTATCCCGTAT
TGACGCCGGGCAAGAGCAACTCGGTCGCCGCATACACTATTCTCAGAATGACTTGGTTGAGTAC
TCACCAGTCACAGAAAAGCATCTTACGGATGGCATGACAGTAAGAGAATTATGCAGTGCTGCCA
TAACCATGAGTGATAACACTGCGGCCAACTTACTTCTGACAACGATCGGAGGACCGAAGGAGCT
AACCGCTTTTTTGCACAACATGGGGGATCATGTAACTCGCCTTGATCGTTGGGAACCGGAGCTG
AATGAAGCCATACCAAACGACGAGCGTGACACCACGATGCCTGTAGCAATGGCAACAACGTTGC
GCAAACTATTAACTGGCGAACTACTTACTCTAGCTTCCCGGCAACAATTAATAGACTGGATGGA
GGCGGATAAAGTTGCAGGACCACTTCTGCGCTCGGCCCTTCCGGCTGGCTGGTTTATTGCTGAT
AAATCTGGAGCCGGTGAGCGTGGCTCTCGCGGTATCATTGCAGCACTGGGGCCAGATGGTAAGC
CCTCCCGTATCGTAGTTATCTACACGACGGGGAGTCAGGCAACTATGGATGAACGAAATAGACA
```

GATCGCTGAGATAGGTGCCTCACTGATTAAGCATTGGTAACTGTCAGACCAAGTTTACTCATAT
ATACTTTAGATTGATTTAAAACTTCATTTTTAATTTAAAAGGATCTAGGTGAAGATCCTTTTTG
ATAATCTCATGACCAAAATCCCTTAACGTGAGTTTTCGTTCCACTGAGCGTCAGACCCCGTAGA
AAAGATCAAAGGATCTTC

>pCYC02_CmR
TTGAGATCCTTTTTTTCTGCGCGTAATCTGCTGCTTGCAAACAAAAAAACCACCGCTACCAGCG
GTGGTTTGTTTGCCGGATCAAGAGCTACCAACTCTTTTTCCGAAGGTAACTGGCTTCAGCAGAG
CGCAGATACCAAATACTGTTCTTCTAGTGTAGCCGTAGTTAGGCCACCACTTCAAGAACTCTGT
AGCACCGCCTACATACCTCGCTCTGCTAATCCTGTTACCAGTGGCTGCTGCCAGTGGCGATAAG
TCGTGTCTTACCGGGTTGGACTCAAGACGATAGTTACCGGATAAGGCGCAGCGGTCGGGCTGAA
CGGGGGGTTCGTGCACACAGCCCAGCTTGGAGCGAACGACCTACACCGAACTGAGATACCTACA
GCGTGAGCTATGAGAAAGCGCCACGCTTCCCGAAGGGAGAAAGGCGGACAGGTATCCGGTAAGC
GGCAGGGTCGGAACAGGAGAGCGCACGAGGGAGCTTCCAGGGGGAAACGCCTGGTATCTTTATA
GTCCTGTCGGGTTTCGCCACCTCTGACTTGAGCGTCGATTTTTGTGATGCTCGTCAGGGGGGCG
GAGCCTATGGAAAAACGCCAGCAACGCGGCCTTTTTACGGTTCCTGGCCTTTTGCTGGCCTTTT
GCTCACATGTTCTTTCCTGCGTTATCCCCTGATTCTGTGGATAACCGTATTACCGCCTTTGAGT
GAGCTGATACCGCTCGCCGCAGCCGAACGACCGAGCGCAGCGAGTCAGTGAGCGAGGAAGCGGA
AGAGCGCCCAATACGCAAACCGCCTCTCCCCGCGCGTTGGCCGATTCATTAATGCAGCTGGCAC
GACAGGTTTCCCGACTGGAAAGCGGGCAGTGAGCGCAACGCAATTAATGTGAGTTAGCTCACTC
ATTAGGCACCCCAGGCTTTACACTTTATGCTTCCGGCTCGTATGTTGTGTGGAATTGTGAGCGG
ATAACAATTTCACACAGGAAACAGCTATGACCATGATTACGCCAAGCTCCTGGAGACCAGCATG
GCTTGCAGTGGCGTTGGTCTCCGGCAACGCGTATATCTGGCCCGTACATCGCGAAGCAGCGCAA
AACGCCTAACCCTAAGCAGATTCTTCATGCAATTGTCGGTCAAGCCTTGCCTTGTTGTAGCTTA
AATTTTGCTCGCGCACTACTCAGCGACCTCCAACACACAAGCAGGGAGCAGATACTGGCTTAAC
TATGCGGCATCAGAGCAGATTGTACTGAGAGTGCACCATAGGGGATCGGGAGATCTCCCGATCC
GTCGACGTCAGGTGGCACTTTTCGGGGAAATGTGTGATCGGCACGTAAGAGGTTCCAACTTTCA
CCATAATGAAATAAGATCACTACCGGGCGTATTTTTTGAGTTATCGAGATTTTCAGGAGCTAAG
GAAGCTAAAATGGAGAAAAAAATCACTGGATATACCACCGTTGATATATCCCAATGGCATCGTA
AAGAACATTTTGAGGCATTTCAGTCAGTTGCTCAATGTACCTATAACCAGACCGTTCAGCTGGA
TATTACGGCCTTTTTAAAGACCGTAAAGAAAAATAAGCACAAGTTTTATCCGGCCTTTATTCAC
ATTCTTGCCCGCCTGATGAATGCTCATCCGGAATTCCGTATGGCAATGAAAGACGGTGAGCTGG
TGATATGGGATAGTGTTCACCCTTGTTACACCGTTTTCCATGAGCAAACTGAAACGTTTTCATC
GCTCTGGAGTGAATACCACGACGATTTCCGGCAGTTTCTACACATATATTCGCAAGATGTGGCG
TGTTACGGTGAAAACCTGGCCTATTTCCCTAAAGGGTTTATTGAGAATATGTTTTTCGTCTCAG
CCAATCCCTGGGTGAGTTTCACCAGTTTTGATTTAAACGTGGCCAATATGGACAACTTCTTCGC
CCCCGTTTTCACCATGGGCAAATATTATACGCAAGGCGACAAGGTGCTGATGCCGCTGGCGATT
CAGGTTCATCATGCCGTTTGTGATGGCTTCCATGTCGGCAGAATGCTTAATGAATTACAACAGT
ACTGCGATGAGTGGCAGGGCGGGGCGTAACTGTCAGACCAAGTTTACTCATATATACTTTAGAT
TGATTTAAAACTTCATTTTTAATTTAAAAGGATCTAGGTGAAGATCCTTTTTGATAATCTCATG

ACCAAAATCCCTTAACGTGAGTTTTCGTTCCACTGAGCGTCAGACCCCGTAGAAAAGATCAAAG
GATCTTC

>pCYC03_KanR
TTGAGATCCTTTTTTTCTGCGCGTAATCTGCTGCTTGCAAACAAAAAAACCACCGCTACCAGCG
GTGGTTTGTTTGCCGGATCAAGAGCTACCAACTCTTTTTCCGAAGGTAACTGGCTTCAGCAGAG
CGCAGATACCAAATACTGTTCTTCTAGTGTAGCCGTAGTTAGGCCACCACTTCAAGAACTCTGT
AGCACCGCCTACATACCTCGCTCTGCTAATCCTGTTACCAGTGGCTGCTGCCAGTGGCGATAAG
TCGTGTCTTACCGGGTTGGACTCAAGACGATAGTTACCGGATAAGGCGCAGCGGTCGGGCTGAA
CGGGGGGTTCGTGCACACAGCCCAGCTTGGAGCGAACGACCTACACCGAACTGAGATACCTACA
GCGTGAGCTATGAGAAAGCGCCACGCTTCCCGAAGGGAGAAAGGCGGACAGGTATCCGGTAAGC
GGCAGGGTCGGAACAGGAGAGCGCACGAGGGAGCTTCCAGGGGGAAACGCCTGGTATCTTTATA
GTCCTGTCGGGTTTCGCCACCTCTGACTTGAGCGTCGATTTTTGTGATGCTCGTCAGGGGGGCG
GAGCCTATGGAAAAACGCCAGCAACGCGGCCTTTTTACGGTTCCTGGCCTTTTGCTGGCCTTTT
GCTCACATGTTCTTTCCTGCGTTATCCCCTGATTCTGTGGATAACCGTATTACCGCCTTTGAGT
GAGCTGATACCGCTCGCCGCAGCCGAACGACCGAGCGCAGCGAGTCAGTGAGCGAGGAAGCGGA
AGAGCGCCCAATACGCAAACCGCCTCTCCCCGCGCGTTGGCCGATTCATTAATGCAGCTGGCAC
GACAGGTTTCCCGACTGGAAAGCGGGCAGTGAGCGCAACGCAATTAATGTGAGTTAGCTCACTC
ATTAGGCACCCCAGGCTTTACACTTTATGCTTCCGGCTCGTATGTTGTGTGGAATTGTGAGCGG
ATAACAATTTCACACAGGAAACAGCTATGACCATGATTACGCCAAGCTCCTGGAGACCAGCATG
GCTTGCAGTGGCGTTGGTCTCCGGCAACGCGTATATCTGGCCCGTACATCGCGAAGCAGCGCAA
AACGCCTAACCCTAAGCAGATTCTTCATGCAATTGTCGGTCAAGCCTTGCCTTGTTGTAGCTTA
AATTTTGCTCGCGCACTACTCAGCGACCTCCAACACACAAGCAGGGAGCAGATACTGGCTTAAC
TATGCGGCATCAGAGCAGATTGTACTGAGAGTGCACCATAGGGGATCGGGAGATCTCCCGATCC
GTCGACGTCAGGTGGCACTTTTCGGGGAAATGTGAGTGGCCAATTTTGCGGCCGCCCTATTTGT
TTATTTTTCTAAATACATTCAAATATGTATCCGCTCATGAGACAATAACCCTGATAAATGCTTC
AATAATATTGAAAAAGGAAGAGTATGAGCCATATTCAACGGGAAACGTCTTGCTCTAGGCCGCG
ATTAAATTCCAACATGGATGCTGATTTATATGGGTATAAATGGGCTCGCGATAATGTCGGGCAA
TCAGGTGCGACAATCTATCGATTGTATGGGAAGCCCGATGCGCCAGAGTTGTTTCTGAAACATG
GCAAAGGTAGCGTTGCCAATGATGTTACAGATGAGATGGTCAGACTAAACTGGCTGACGGAATT
TATGCCTCTTCCGACCATCAAGCATTTTATCCGTACTCCTGATGATGCATGGTTACTCACCACT
GCGATCCCTGGGAAAACAGCATTCCAGGTATTAGAAGAATATCCTGATTCAGGTGAAAATATTG
TTGATGCGCTGGCAGTGTTCCTGCGCCGGTTGCATTCGATTCCTGTTTGTAATTGTCCTTTTAA
CAGCGATCGCGTATTTCGTCTCGCTCAGGCGCAATCACGAATGAATAACGGTTTGGTTGATGCG
AGTGATTTTGATGACGAGCGTAATGGCTGGCCTGTTGAACAAGTCTGGAAAGAAATGCATAAAC
TTTTGCCATTCTCACCGGATTCAGTCGTCACTCATGGTGATTTCTCACTTGATAACCTTATTTT
TGACGAGGGGAAATTAATAGGTTGTATTGATGTTGGACGAGTCGGAATCGCAGACCGATACCAG
GATCTTGCCATCCTATGGAACTGCCTCGGTGAGTTTTCTCCTTCATTACAGAAACGGCTTTTTC
AAAAATATGGTATTGATAATCCTGATATGAATAAATTGCAGTTTCATTTGATGCTCGATGAGTT
TTTCTAACTGTCAGACCAAGTTTACTCATATATACTTTAGATTGATTTAAAACTTCATTTTTAA

TTTAAAAGGATCTAGGTGAAGATCCTTTTTGATAATCTCATGACCAAAATCCCTTAACGTGAGT
TTTCGTTCCACTGAGCGTCAGACCCCGTAGAAAAGATCAAAGGATCTTC
